# Sensitivity to photoperiod is a complex trait in *Camelina sativa*

**DOI:** 10.1101/2024.10.25.620367

**Authors:** Bryan A Ramirez-Corona, Erin Seagren, Carissa Sherman, Takato Imaizumi, Christine Queitsch, Josh Cuperus

## Abstract

Day neutrality, or insensitivity to photoperiod (day length), is an important domestication trait in many crop species. Although the oilseed crop *Camelina sativa* has been cultivated since the Neolithic era, day-neutral accessions have yet to be described. We sought to leverage genetic diversity in existing germplasms to identify *C. sativa* accessions with low photoperiod sensitivity for future engineering of this trait. We quantified variation in the photoperiod response across 161 accessions of *C. sativa* by measuring hypocotyl length of four-day-old seedlings grown in long-day and short-day conditions, finding wide variation in photoperiod response. Similarly, soil-grown adult plants from selected accessions showed variation in photoperiod response in several traits; however, photoperiod responses in seedling and adult traits were not correlated, suggesting complex mechanistic underpinnings. Although RNA-seq experiments of the reference accession Licalla identified several differentially regulated *Arabidopsis* syntelogs involved in photoperiod response, including *COL2, FT, LHY* and *WOX4*, expression of these genes in the accessions did not correlate with differences in their photoperiod sensitivity. Taken together, we show that all tested accessions show some degree of photoperiod response, and that this trait is likely complex, involving several and separable seedling and adult traits.

**Significance Statement:** Day neutrality (photoperiod insensitivity) is a common trait in domesticated crops; however, the ancient oilseed crop *Camelina sativa* has remained photoperiod-sensitive, which likely limits seed yields. Here, we show that photoperiod sensitivity is conserved across many *C. sativa* cultivars, albeit to different degrees, and we establish that photoperiod sensitivity is a complex trait, which will require genetic engineering to achieve day neutrality.

## Introduction

Climate change, population growth and the loss of arable land are major challenges that threaten food security. One approach to ensuring food security is the development of sustainable low-resource crops that can grow on marginal land, are stress-resistant and are high yielding (Berti *et al*., 2016). One such crop is *Camelina sativa*, a low-resource oilseed crop that is amenable to genetic engineering (Berti *et al*., 2016). Because of *C. sativa’s* agricultural potential, recent studies have developed genetic resources, genome sequence and expression data, among other resources for this crop (Kagale *et al*., *2016*, King *et al*., 2019, Luo *et al*., 2019, Gomez-Cano *et al*., 2020).

*C. sativa* grows well in marginal soils, adapts readily to adverse environmental conditions, and has low water and nutrient requirements compared to other oilseed crops (Vollmann & Eynck 2015). Unlike the high-yielding oilseed crop *Brassica napus* (canola), *C. sativa* is resistant to common *Brassicaceae* pests and pathogens (Séguin-Swartz *et al*.,2009). Camelina seed oil content ranges from 36-47% by weight, with exceptionally high levels of essential and omega-3 fatty acids, a profile broadly useful in food, animal feed, industrial bioproducts and biofuel (Berti *et al*., 2016, Yuan & Li 2020). In field trials, the crop reduces weed biomass through the release of inhibitory chemical compounds, demonstrating its allelopathic potential (Ghidoli *et al*., 2023). Camelina is readily transformable using the floral dip method, enabling genetic engineering (Lu & Kang 2008). Its short life span (85-100 days) and ability to be planted and harvested using conventional equipment make field trials of engineered plants straightforward (Malik *et al*.,2018).

The major reason for *C. sativa’s* displacement by canola is the crop’s modest seed yield (Obour *et al*., 2015, Berti *et al*., 2016). In many domesticated crops such as canola, rice, maize, sorghum, potato and others, the loss of the photoperiod response – the acquisition of day neutrality – is common and has allowed farming of these crops at higher latitudes (Doebly *et al*., 2016). As a long-day plant, *C. sativa* flowers in the spring at higher latitudes, thereby accumulating comparatively little vegetative biomass to produce carbohydrates and ultimately seeds. In canola, vegetative biomass is the primary determinant of seed yield (Bennet *et al*. 2017, Zhang & Flottman 2017, Chen *et al*. 2021), so it stands to reason that one way to increase *C. sativa* seed yield is to generate day-neutral varieties. Thus, to engineer or breed *C. sativa* cultivars with higher seed yield, it is imperative to understand both the extent of natural variation of the photoperiod response in this crop and its genetic underpinnings.

Here, we quantified the photoperiod response across 161 diverse *C. sativa* accessions by recording hypocotyl length and germination rate of seedlings grown in long-day (LD) and short-day (SD) conditions. Using Licalla as a reference accession, we categorized accessions as either low- or high-responsive to photoperiod. Eight accessions with low, medium, or high photoperiod sensitivity on hypocotyl growth were selected for measurements of adult developmental traits associated with photoperiod responses. Seedling photoperiod responses were not predictive of photoperiod responses in adult agronomic traits. Gene expression levels of four photoperiod response genes – *COL2, FT, LHY* and *WOX4* – did not explain the accession-specific differences. In sum, *C. sativa* accessions show a range of photoperiod sensitivity; however, there is little correlation across different traits associated with photoperiod response, suggesting complex mechanistic underpinnings.

## RESULTS

To quantify the phenotypic variation in photoperiod response, we grew 161 *C. sativa* accessions (Supplemental Table 1) under LD or SD conditions and measured the hypocotyl length of four-day-old seedlings (Supplemental Table 1). Accessions were split into 13 experimental batches. Each batch consisted of 32 seedlings per photoperiod treatment for each accession. The reference accession Licalla (Gehringer *et al*., 2007) included in each batch (Experimental Procedures). While the accessions varied greatly in the number of seeds that germinated on the first day, 73% of LD-grown seeds and 75% of SD-grown seeds did so, with a mean germination rate per accession of 85% (Supplemental Figure 1, Supplemental Table 3). To account for differences in hypocotyl length that were due to delayed germination, we restricted our analysis to seedlings that germinated on the first day and excluded eight accessions from further analysis.

As expected, LD-grown seedlings had generally shorter hypocotyls than SD-grown seedlings (Figure 1 A and B). To quantify differences in photoperiod sensitivity among accessions, we calculated the mean difference (MD) in hypocotyl length for each accession between SD and LD conditions and adjusted for experimental batch effects (Experimental Procedures, Supplemental Figure 2). To correct for batch effects, we included seedlings of the reference accession Licalla on all plates and calculated a normalized mean difference (NMD) in hypocotyl length by dividing each tested accession’s MD value by the MD value of the respective Licalla seedlings (Experimental Procedures). In short, a fully day-neutral accession would show an NMD of zero while Licalla would show an NMD of 1.

**Figure 1:**
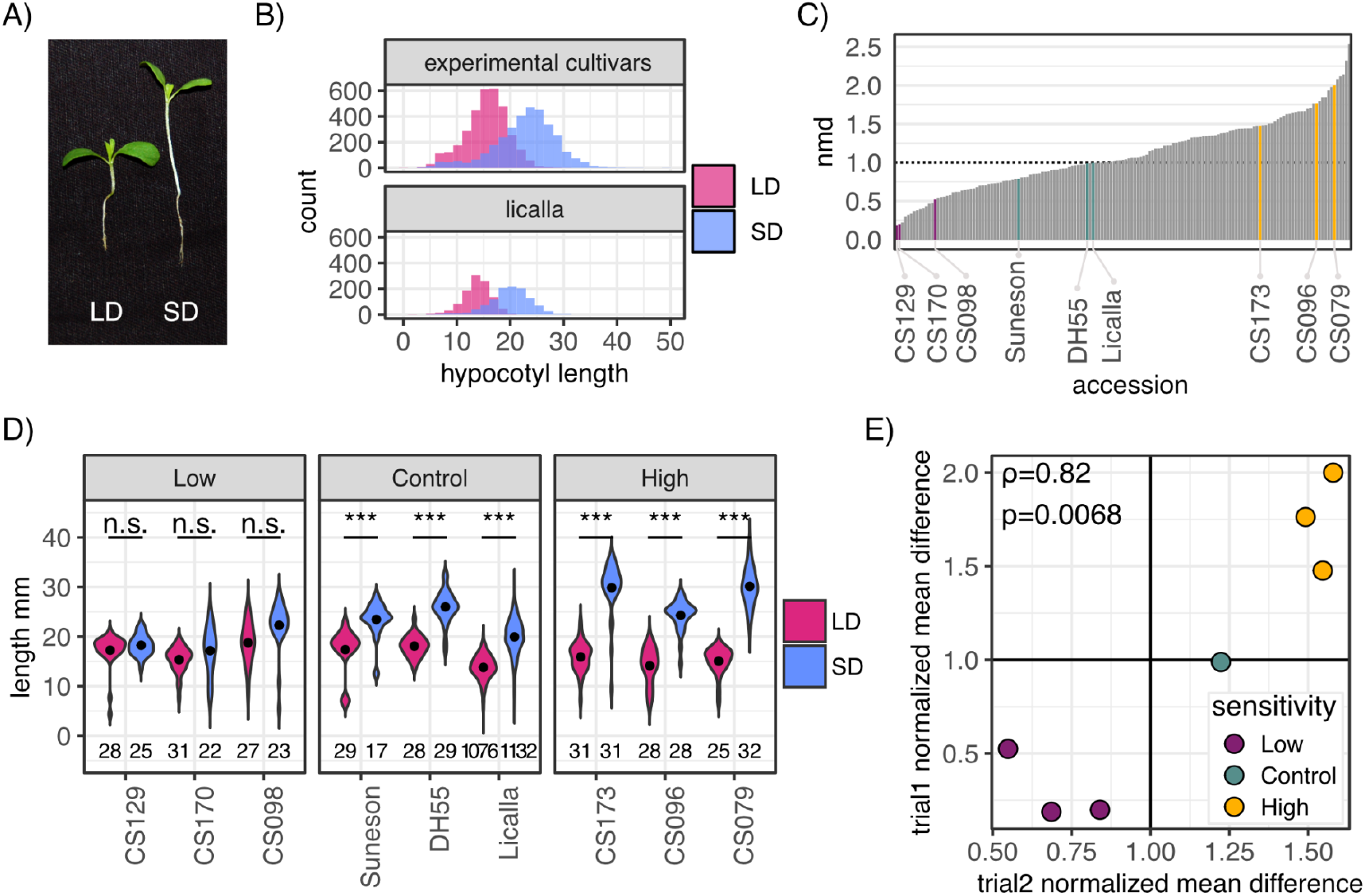
Quantification of seedling photoperiod responses in 161 accessions of *Camelina sativa*. A) Representative image of *C. sativa* seedlings (Licalla accession) grown under long-day (LD) or short-day (SD) conditions. B) Distribution of hypocotyl lengths from four-day-old seeds germinated on day one for all experimental accessions and Licalla. Long photoperiod treatment (magenta) yields shorter hypocotyls than short photoperiod (blue) treatment. C) Normalized mean difference (NMD) for all accessions corrected by batch. Low photoperiod-sensitive accessions (CS170, CS129, CS098) are marked in purple, high photoperiod-sensitive accessions (CS173, CS096 and CS079) are marked in gold and control accessions Licalla (CS002), DH55 and Suneson (CS001) are marked in green. All accessions show some degree of photoperiod response. D) LD and SD hypocotyl lengths of selected accessions in both SD and LD treatments, X axis is ordered by increasing NMD (two sided t-test, *: p≤ 0.01, **: p≤ 0.001, ***: p≤ 0.0001). Low photoperiod-sensitive accessions do not show significant differences in the mean hypocotyl length between photoperiod treatments. E) Correlation of NMD values between two trials of hypocotyl assays shows that exact rank order is not preserved but the differences between low and high photoperiod-sensitive accession remain. Trials show significant positive correlation with each other (Spearman rank correlation, ρ=0.82, p-value=0.0068).

*C. sativa* accessions varied greatly in NMD, with the well-known accessions DH55 (Kagale *et al*. 2014) and Suneson (CS001, Li *et al*. 2021) being less photoperiod sensitive than Licalla; overall, 86 accessions were more photoperiod sensitive (NMD>1) than Licalla while 66 were less photoperiod sensitive (NMD<1) based on photoperiodic hypocotyl growth phenotypes. Most accessions showed significant differences in hypocotyl length between LD and SD (n=121), including Licalla, DH55 and Suneson (Supplemental Figure 3). Next, we categorized accessions as either low or high photoperiod-sensitive. Accessions with an NMD less than 1 and without significant differences in hypocotyl length between LD and SD, were categorized as low photoperiod-sensitive (n=19). Accessions with an NMD greater than 1 and with significant differences in hypocotyl length between LD and SD were categorized as high photoperiod-sensitive (n=76). The remaining accessions were excluded from further analyses.

While hypocotyl measurements in seedlings makes it feasible to test many accessions efficiently, we next addressed how well these early-stage differences in photoperiod response were maintained in adult plants. In *Arabidopsis*, photoperiods affect hypocotyl growth and flowering time response, and mutants affecting circadian time measurement often display hypocotyl and flowering phenotypes (Niwa *et al*., 2009, Nagel & Kay 2012). Therefore, we speculated that accessions with high or low photoperiod sensitivity may be caused by differences in seasonal time measurement ability. If so, these seedlings might show altered photoperiodic phenotypes as adult plants. To test this hypothesis, we selected a subset of accessions at both ends of the photoperiod response spectrum. Because differences in NMD can be the result of low or variable germination rates and low accession health, we manually screened images of candidate accessions for high germination rates and robust seedling growth in both LD and SD conditions (Supplemental Figure 4). We selected three high photoperiod-sensitive accessions (CS173, CS096 and CS079) and three low photoperiod-sensitive accessions (CS170, CS129 and CS098) that fit these criteria (Figure 1D), in addition to the control accessions DH55 and Licalla.

While the selected accessions were grown to adulthood, we conducted a validation experiment that tested the hypocotyl photoperiod response of these accessions in a single experiment with twice the number of seedlings per test condition. The majority of seeds germinated on the first day, allowing us to focus our analysis on day-one seedlings (Supplemental Figure 5A). Two of the three low photoperiod-sensitive accessions did not show significant differences in hypocotyl lengths between LD and SD conditions (Supplemental Figure 5, CS170: p-value=0.11, CS098: p=0.23). However, a third accession, CS129, now showed a significant difference between LD and SD hypocotyl length (p-value=0.002) not observed in the first trial, likely due to the increased power in the validation experiment. As expected from the previous trial, both the control and high photoperiod-sensitive accessions showed significant differences in hypocotyl length between LD and SD. NMD measurements between the two trials were significantly correlated (Figure 1E, Spearman’s ρ=0.82, p-value=0.0068). Although the exact rank order of photoperiod sensitivity was not preserved between the two trials, the overall separation of high photoperiod-sensitive and low photoperiod-sensitive accessions was validated.

We next asked whether these differences were maintained in adult traits. Height, root mass, flowering time and seed yield are important agronomic traits for breeding and crop development and were quantified in adult plants. We grew ten plants of each selected accession under LD and SD conditions. Four plants from each accession were used for gene expression measurements and root mass measurements at day 20 (Experimental Procedures). The remaining six plants of each accession were grown for 65 days before drying for seed harvesting (Experimental Procedures). Contrary to our observations in the hypocotyl assay, the central stalk of soil-grown plants in SD conditions were shorter in height than plants in LD conditions, consistent with reduced vegetative growth (Figure 2A). Considering the mean difference in adult plant height, the initial separation of photoperiod sensitivity observed in the hypocotyl assay was lost as early as in the third week of plant growth and remained so until the end of the trial (Supplemental Figure 6). Neither rank order nor the categories of high and low photoperiod-sensitive groups were preserved.

**Figure 2:**
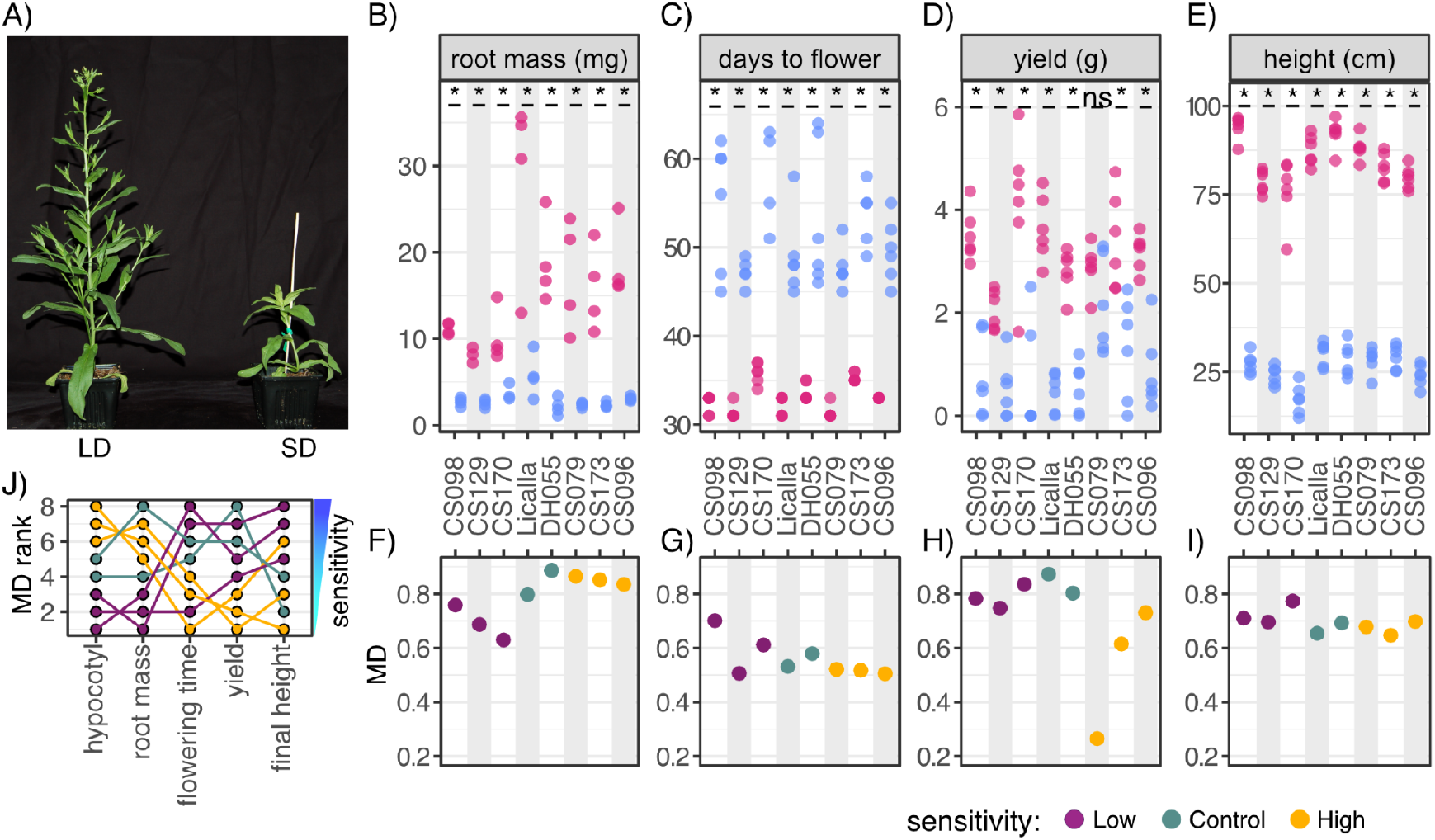
Rank order of photoperiod sensitivity is not maintained across traits. A) Representative image of 5-week-old soil-grown Camelina plants in LD (left) and SD (right) conditions. B-E) Pink points indicate LD treatment and blue points indicate SD treatment. X-axis shows accessions ordered left to right by increasing hypocotyl length NMD. Y-axis shows the measured adult trait listed at the top of the graph. B) Root mass measured in 20-day-old plants with significant differences between SD and LD conditions labeled (two sided t-test, * < p-value 0.05). C) All accessions showed significant differences in the number of days to flowering between SD and LD conditions (top, two sided t-test, * < p-value 0.05). D) All accessions except CS079 showed significant differences in yield (total seed weight in g) between SD and LD conditions (two sided t-test, *< p-value 0.05). E) All accessions showed significant differences in the height of the central stalk (cm) at 41 days, between SD and LD conditions (two sided t-test, *< p-value 0.05). F-I) X-axis shows accessions ordered left to right by increasing hypocotyl length NMD. Purple points indicate accessions with low photoperiod sensitivity in hypocotyls, yellow points indicate accessions with high photoperiod sensitivity in hypocotyls and green points indicate control accessions. Y-axis shows the corresponding MD value for each accession for corresponding trait in the graph above. J) Ranked photoperiod response for the eight accessions across adult traits using MD for each trait.

Root mass at day 20, days to flowering and seed yield were also quantified for each accession. For both root mass and days to flowering, all accessions showed significant differences between LD and SD conditions (Figure 2B-E). Similarly, the total seed weight for each accession significantly increased in LD conditions, except for the CS079 accession for which there was no difference (Figure 2D). This observation is notable because this accession showed high photoperiod sensitivity in the seedling trait hypocotyl length, but it is the least photoperiod-sensitive accession for the adult plant traits total seed weight. A plant’s seed yield is most readily measured as total seed weight, however, seed weight divided by the total number of seeds produced is also highly informative for breeders. Here, we approximated this measure by weighing 100 seeds for each tested accession (individual seed weight). The average individual seed weight and estimated number of seeds were significantly correlated in the control and low photoperiod-sensitive accessions, indicating that these accessions achieve higher total seed weight in LD by increasing both the total number of seeds and the individual seed weight (Supplemental Figure 7A). High photoperiod-sensitive accessions, however, failed to show strong correlation between total seed number and average individual seed weight. Thus, the higher total seed weight in these accessions in LD conditions was the result of additional seeds of similar size that were markedly smaller than those in low photoperiod-sensitive accessions.

For each measured trait, we calculated the MD of each accession (Figure 2F-I), normalized by the Licalla MD value. Neither rank order nor the observed separation of low and high photoperiod-sensitive accessions was maintained between our hypocotyl assay and the adult traits (Figure 2J). Spearman rank correlations of trait MD values showed no significant correlations between any of the adult traits (Supplemental Figure 8). Due to this lack of concordance in MD rank across traits, none of the accessions can be singularly classified as less photoperiod-sensitive than the others. This result is consistent with photoperiod response being a complex trait in this crop.

Although much is known in *Arabidopsis thaliana* about photoperiod response and the genes that regulate it (Nagel & Kay 2012, Song *et al*., 2018), far less is known about how the *C. sativa* syntelogs behave and their utility as potential markers for phenotypic traits of interest. To address this knowledge gap, we performed bulk RNA sequencing on the aerial tissue of 3-week old Licalla plants grown in either LD or SD conditions (Experimental Procedures). Of the detected 40,468 genes, 218 were found to be differentially expressed between LD and SD conditions. Specifically, 151 genes were upregulated, and 67 genes were downregulated in LD conditions, relative to SD (Figure 3 A). Of these genes,126 had known Arabidopsis orthologs, 98 were upregulated and 28 were downregulated. Of the upregulated genes, four genes were of particular interest: *CONSTANS-LIKE 2* (COL2), *FLOWERING LOCUS T* (*FT*)/*TWIN SISTER OF FT* (*TSF*), *LATE ELONGATED HYPOCOTYL* (*LHY*) and *WUSCHEL RELATED HOMEOBOX 4* (*WOX4*), syntelogs of *Arabidopsis* that are involved in photoperiod-controlled developmental responses. *COL2* is a zinc finger protein with sequence similarity to the flowering gene *CONSTANS* and has been implicated in flowering time regulation in other plants (Ledger 2001, Liu 2021, Liang 2023). *FT* is a florigen that, along with *TSF*, acts as a mobile signal to induce the vegetative to flowering transition (Song *et al*., 2015, Wang *et al., 2020*, Lee *et al*., 2023); both are syntelogs of the highly differentially expressed *C. sativa* gene Csa05g068740, which had nearly an 8-fold change in expression between LD and SD. *LHY* is a core circadian clock gene that in *Arabidopsis* is involved in the regulation of several developmental processes including flowering time and the *FT* locus (Fujiwara *et al*., *2008*, Nagel & Kay 2012). While *WOX4* in *Arabidopsis* is primarily involved in cell division and vascular proliferation, several WOX transcription factors are involved in floral development (Costanzo *et al*., 2014). Other syntelogs of flowering time regulators, such as *EARLY FLOWERING 3* (ELF3), *GIGANTEA* (GI) and *CONSTANS* (CO), were not differentially expressed in our data (Supplemental Figure 9, Nagel & Kay 2012). Although these selected examples had syntelogs in *Arabidopsis*, many other differentially expressed genes did not (n=92) suggesting there are many more potential genes of interest to dissect.

**Figure 3:**
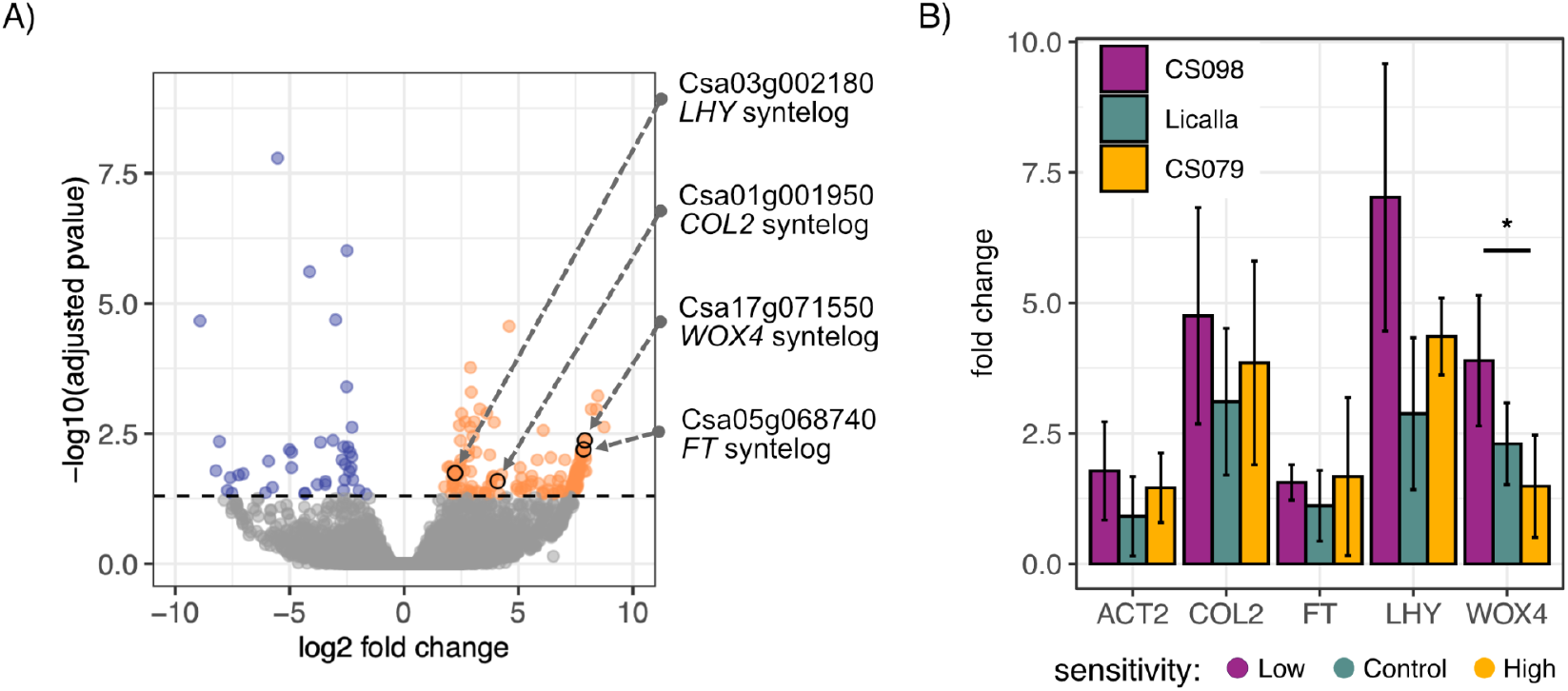
Expression of known photoperiod response genes is higher in LD but no differences are observed among accessions with low and high photoperiod sensitivity in seedlings. A) Expression differences in LD 3-week-old Licalla leaves relative to SD. A total of 40,468 genes were detected with 151 upregulated and 67 down regulated. Syntelogs for potential photoperiod responsive genes *COL2, LHY, WOX4* and *FT* are highlighted. B) qPCR of the low photoperiod-sensitive accession CS098 (purple), the control accession Licalla (green), and the high photoperiod-sensitive accession CS079 (gold). Expression of selected genes is higher in LD conditions. Expression of *WOX4* differs significantly between CS098 and CS079 accessions (two sided t-test, p=0.049, Bonferroni corrected); the remaining genes did not show significant expression differences. Columns show the mean fold change and error bars show standard deviation.

Having identified potential photoperiod response regulators upregulated in LD in Licalla plants, we asked if expression differences of these genes could explain the differences observed between accessions in adult traits. We extracted RNA from aerial tissue of 20-day old plants grown in either LD or SD, using one genotype from the low sensitivity (CS098), high sensitivity (CS079) and reference (Licalla) groups. We performed qPCR using primers for *COL2, FT, LHY, WOX4, ACT2* and *SEC3* (Supplemental Table 2). *ACT2* was included as a control and SEC3 was used as the calibrator for calculating fold change (2^−ΔΔCq^) relative to SD treatment (Supplemental data 3, Chau *et al*., *2018*). All selected genes were upregulated in LD conditions. The only significant accession differences observed were in *WOX4* between the high and low sensitivity lines (Figure 4 B). We noted that in particular for *FT* and *WOX4*, the magnitude of expression differences between the RNA-sequencing results and the PCR data was large (230-fold vs. 2-fold). We suspect that this difference is due to the difference in the time of tissue collection (RNA-seq ZT4; qPCR, ZT8). Nevertheless, the expression patterns of selected genes were insufficient to tease apart the observed phenotypic differences among these accessions. Finding genes whose expression patterns better explains the phenotypic variation observed in Camelina photoperiod response will be necessary for future studies.

## Discussion

Here, we quantified differences in photoperiod response in a panel of 161 *Camelina sativa* accessions, observing a wide spectrum of photoperiod sensitivity in the seedling trait hypocotyl length. Several accessions showed little difference in hypocotyl length between SD and LD conditions, appearing near day-neutral in this early trait. Accessions were categorized as either low or high photoperiod-sensitive, and three accessions at both ends of the phenotypic spectrum were selected for testing photoperiod response in adult traits. None of the selected accessions were found to be day neutral at the adult stage. We observed significant differences between LD and SD treatments in the adult traits of root mass, height, flowering time, and seed yield. However, there was no meaningful correlation between photoperiod sensitivity in the seedling trait hypocotyl length and photoperiod sensitivity in the measured adult traits. In fact, rank order of photoperiod sensitivity differed across all the measured adult traits, suggesting photoperiodic phenotypes of the accessions we selected have altered responses in each photoperiodic response, but seasonal time measurement mechanisms were not altered.

Our results suggest that day neutrality may not be present among existing *C. sativa* germplasm. If so, breeders cannot rely on introgression of day-neutrality from existing accessions for the development of high-yielding, day-neutral *C. sativa*. Rather, genetic engineering of photoperiod measurement mechanisms will be required to generate such lines. In order to facilitate the identification of possible engineering targets, we conducted RNA-seq with the reference accession Licalla. Of the 151 significantly upregulated genes, we selected four *Arabidopsis* syntelogs involved in photoperiod response, *COL2, FT, LHY* and *WOX4*, for expression studies in low and high photoperiod-sensitive accessions. However none of the genes showed accession-specific differences in expression. Other known *Arabidopsis* syntelogs, *CO, ELF3* and *GI*, were detected in our data set but were not found to be differentially expressed between LD and SD, however, these genes are known to peak in the evening, whereas our data was collected in the morning (ZT4).

The molecular basis of day neutrality in other crops has been shown to be complex (Lin *et al*., 2021, Wang *et al*., 2023). Studies in rice, tomato barley and canola, among other crops have uncovered some of the genes and candidate loci involved in reducing photoperiod response and increasing yields (Turner *et al*., 2005, Comadran *et al*., 2012, Wang *et al*., 2016, Soyk *et al*. 2017, Wei *et al*., 2017, Liu *et al*., 2018, Zhang *et al*., 2018, Lu *et al*., *2019*, Song *et al*., 2020, Zong *et al*., 2021). In barley, these studies have yielded a complex picture with different alleles of *TERMINAL FLOWER1*/*CENTRORADIALIS* conferring an advantage under different environmental conditions (Comadran *et al*., 2012). In spring-sown barley, a Pseudo-response regulator *Ppd-H1* variant delays flowering specifically in long days, illustrating that variation in diverse genes associated with clock function and photoreception can confer a weaker photoperiod response (Turner *et al*., 2005). In canola, the world’s second most important oilseed crop, several dozen loci contribute to variation in flowering time among cultivars, consistent with the crops complex allotetraploid nature (Schiessl 2020, Song *et al*., 2020). In tomato, domesticated day-neutral lines have been found to have altered circadian rhythms that appear to confer fitness under long-day conditions (Müller *et. al* 2015) In several crops including tomato and rice variations in regulatory DNA and changes in promoter enhancer interaction are implicated in the acquisition of day neutrality (Takahashi *et al*., 2009, Zhang *et al*., 2018). An attenuated photoperiod response is often associated with the loss for the vernalization requirement-the need for a ‘winter’ period before flowering (Malik *et al*., 2018).

Without the benefit of *C. sativa* varieties with stark and consistent differences in photoperiod sensitivity, engineering this trait will be a formidable challenge. A first step toward disentangling the genetic underpinnings of photoperiod sensitivity in *C. sativa* would be detailed expression studies across development and tissues to shed light on genes that consistently show photoperiod-sensitivity. Our simple expression experiment discovered 92 differentially expressed genes without *A. thaliana* syntelogs and is a well-suited starting point for such future investigations. Additionally, it will be necessary to identify traits or sets of traits that are most predictive of day neutrality to facilitate the engineering and breeding of *C. sativa* varieties that combine day-neutrality and high seed yields with the crop’s other favorable agronomic properties.

## Experimental Procedures

### Accessions / Plant Materials & Growth Conditions / Camelina Cultivation

*Camelina sativa* stocks consisted of 160 accessions generously provided by Jennifer Lachowiec from Montana State University as well as DH55 from Agriculture Agri Food Canada (Kagale *et al*., 2014, Li *et al*., 2021). All accessions were seeded in soil (Sunshine Mix #4) and grown in one of two photoperiod conditions, LD (16 h light 8 h dark; 250 µmol m^−2^ sec^−1^; R:FR ratio=1) or SD (8 h light 16 h dark; 500 µmol m^−2^ sec^−1^; R:FR ratio=1) at 22°C. Valoya BX LEDs lights were used. For seed collection, plants were grown for ∼ 9 weeks before water supply was slowly reduced to dry plants for harvesting. Seeds from plants grown under SD and LD conditions were combined into one seed stock per accession which was used for all subsequent experiments.

Seeds were stored in coin envelopes under open air conditions or in closed plastic containers containing desiccants (DRIERITE anhydrous calcium sulfate).

### Hypocotyl Elongation Assay

To characterize photoperiod-dependent hypocotyl elongation differences, 161 *C. sativa* accessions were assayed under SD and LD conditions. Seeds were sown on clear square grid plates (Genesee Cat# 26-275) containing Murashige and Skoog (MS) (PhytoTechnology Laboratories) agar media (1x MS basal salts, 1x MS vitamin powder, 1% sucrose, 0.3% phytagel, 0.5 g/L MES hydrate). Seeds were sterilized by 10-minute exposure to 70% ethanol and 0.5% Tween 20 (ThermoFisher Scientific) followed by 5-minute exposure to 95% ethanol while being shaken vigorously. Sterile seeds were suspended in 0.1% agarose and 8 seeds were pipetted onto each experimental plate. After plating, seeds were placed in the dark at 4°C for ∼24 hours to synchronize germination. The experimental plates were then split between LD growth conditions (16 h light 8 h dark; 100 µmol m^−2^ sec^−1^) or SD growth conditions (8 h light 16 h dark; 125 µmol m^−2^ sec^−1^). For each accession, 4 plates (32 seeds) were run per photoperiod condition, for a total of 64 experimental seeds per accession. Each condition and accession pairing was simultaneously run with one *C. sativa* Licalla plate, which was used as a control due to its known photoperiod-dependent hypocotyl length response.

Following germination synchronization, seed plates were grown for four days in Conviron TC26 growth chambers. During the growth period, each day plate locations within the growth chamber were rotated and germinated seeds were marked. At the end of the growth period, plates were examined under a dissecting microscope and the ends of the hypocotyls were marked prior to imaging.

### Image Analysis

Image analysis was conducted using ImageJ 1.53. Using a graphics tablet with stylus (Wacom Intuos3 PTZ1230), 10 measurements of the plate grid length were taken and averaged to set the pixel to millimeter scale. To measure the hypocotyl length, each hypocotyl was manually traced from end to end using the marks that were made under the microscope as a guide. Contaminated seeds were excluded.

### Hypocotyl Assay Data Analysis

Data collected from the hypocotyl assay was imported into R (3.6.0) for analysis and ggplot2 (3.4.4) was used to generate figures. Only Seeds that germinated on the first day of the growth period and that had a non-zero hypocotyl length were included in this analysis. To test for significant differences in photoperiod-dependent hypocotyl length response within each accession we used a two sided t-test on hypocotyl length measurements between LD and SD. To compare response between accessions, the difference in mean hypocotyl length between LD-grown seedlings and SD-grown seedlings was calculated for each accession and then divided by the LD hypocotyl length of the accession (*MD*_*accession*_). To account for batch effects, we then divided accession MD values by the corresponding MD value of Licalla for the corresponding batch thus calculating normalized mean difference (NMD).

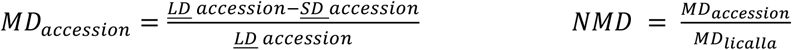

### Soil-Grown Assay

Six selected putative high and low photoperiod sensitivity accessions were grown in soil to determine if the differential photoperiod response observed in hypocotyls was reflected in adult phenotypes. Two accessions were grown as controls: *C. sativa* Licalla whose photoperiod response was already characterized, and DH55, which was used as the *C. sativa* reference genome.

For each accession, 10 plants were grown under LD conditions and 10 were grown under SD conditions as outlined in <u>Growth Conditions</u>. When the plants reached 3 weeks of age, 4 LD-grown and 4 SD-grown plants per accession were selected at random to be harvested. The aerial tissue, including leaves and stems, was separated from the root tissue and flash frozen in liquid nitrogen. Collected tissue was stored at -80°C prior to RNA extraction. The root tissue was washed of soil and debris, and then dried at 80°C for 24 hours prior to weighing.

The remaining 6 plants per accession and photoperiod condition were grown to adulthood. Weekly height measurements were taken from soil level to the top of the central stalk with a meter stick from week 3-6. Each day, plants were checked and the date of emergence of the first flower was recorded for each plant. After approximately 65 days, watering frequency was gradually reduced to allow the plants to dry for harvesting. Seeds were harvested and stored as in previous experiments, however, the seeds of each plant were stored individually and weighed. Additionally, to obtain an average mass per seed, 100 seeds from each plant were weighed.

### RNA extraction

Plant aerial tissue was stored in -80°C. For RNA extraction, tissue was ground with liquid nitrogen using a mortar and pestle and suspended in 10 mL of QIAIzol reagent (Qiagen). This suspension was separated via centrifugation (10 minutes at 4,000g, 4°C) and 5 mL of the supernatant was mixed with 1 mL chloroform. This mixture was centrifuged (15 minutes at 4,000g, 4°C) and the resulting aqueous phase was transferred to a new tube and incubated for 15 minutes at room temperature with 2.5 mL high salt buffer (0.8 M sodium citrate, 1.2 M NaCl) and 2.5 mL isopropanol to precipitate RNA. The precipitate mixture was centrifuged (30 minutes at 4,000g, 4°C) and the resulting pellet was washed with 10 mL of cold 70% ethanol prior to resuspension in 200 µL of RNase free water.

### RNA-seq

RNA sequencing libraries were prepared using extracted RNA and Illumina Stranded mRNA Prep kit. Reference accession Licalla were grown to 3 weeks old and three samples from aerial tissue were prepared per photoperiod treatment at Zeitgeber Time 4 (ZT4, n=6). Sequencing was performed on NextSeq2000. Reads were trimmed using Trim Galore (0.6.10) default settings. Alignment of trimmed reads to the *C. sativa* genome (Kagale *et al*., 2014) was done using STAR (2.7.11.b) default settings and counts were quantified using htseq-counts (2.0.3) using specifications “-m union -r pos -i gene_name -a 10 –stranded=no”. Count data was downloaded into R (3.6.0) and differential expression analysis was conducted using DESeq2 (3.19)

### qPCR

Four samples were collected per accession (CS098, Licalla and CS079) and condition (LD and SD) for qPCR (N=18). Aerial tissue was collected from 3 week-old plants at ZT8 First strand synthesis was done using SuperScript IV first strand synthesis Kit with ezDNase and RNaseH treatment (Invitrogen: 11766050) and cDNA was purified using the Zymo DNA Clean and concentrator kit. RT-qPCR was performed on a CFX Connect Real-Time System (BioRad) using 2x KAPA HiFi HotStart ReadyMix, 0.4X SYBR and 100 mM primers. For amplification, the following program was used: initial denaturation at 98°C for 30 seconds, followed by 40 cycles of 98°C for 20 seconds, 61°C for 30 seconds, and 72°C for 15 seconds. SEC3A was utilized as the calibrator gene for calculating sample ΔCq values. For tested accessions, ΔΔCq was calculated relative to LD and fold change was calculated as 2^−ΔΔCq^. Primers are listed in Supplemental Table 2.

## Accession Numbers

Transcriptomic data can be found in the NIH short read archive (https://submit.ncbi.nlm.nih.gov/subs/sra/) under BioProject ID: PRJNA1086893.

## Acknowledgments

We thank Dr. Jennifer Lachowiec, Montana State University, for Camelina accession seeds, and Agriculture Agri Food Canada for providing DH55 seeds. We thank Stanley Fields for thoughtful discussion and editing of the manuscript. This work was supported by the National Science Foundation (RESEARCH-PGR grant no. 1748843 to S.F. and C.Q. and PlantSynBio grant no. 2240888 to C.Q.). BRC is supported by an NSF Postdoctoral Research Fellowship in the Biology Program (Grant Number 2305660).

## Short Legends for Supporting information

Supplemental Figure 1. Accessions show a range of both germination day and proportion germinated.

Supplemental Figure 2. Mean hypocotyl length significantly correlated with Licalla hypocotyl length when stratified by batch.

Supplemental Figure 3. The majority of accessions show significant differences between SD and LD hypocotyl lengths.

Supplemental Figure 4. Selecting healthy accessions for validation experiments.

Supplemental Figure 5: Germination rate and hypocotyl length of LD and SD plants for selected low photoperiod-sensitive, high photoperiod-sensitive and control accessions.

Supplemental Figure 6. Photoperiod sensitivity in adult plant height varies among accessions over time.

Supplemental Figure 7. Average individual seed weight and estimated seed number are least correlated in high sensitivity accessions.

Supplemental Figure 8: Trait NMD is not significantly correlated across tested Camelina accessions.

Supplemental Figure 9: Common flowering time regulators are not differentially expressed between LD and SD conditions.

Supplemental Table 1: Name and source information for the 161 accessions included in this study

Supplemental Table 2: Primer sequences for qPCR. Supplemental Table 3: Germination day summary

## References

Bennett EJ, Brignell CJ, Carion PWC, Cook SM, Eastmond PJ, Teakle GR, Hammond JP, Love C, King GJ, Roberts JA et al. (2017). “Development of a Statistical Crop Model to Explain the Relationship between Seed Yield and Phenotypic Diversity within the Brassica Napus Gene pool.” Agronomy 7(2).

Berti M, Gesch R, Eynck C, Anderson J, & Cermak S (2016). Camelina uses, genetics, genomics, production, and management. Industrial Crops and Products, 94(2016), 690–710. 10.1016/j.indcrop.2016.09.034

Chao WS, Wang H, Horvath DP, & Anderson JV (2019). Selection of endogenous reference genes for qRT-PCR analysis in Camelina sativa and identification of FLOWERING LOCUS C allele-specific markers to differentiate summer- and winter-biotypes. Industrial Crops and Products, 129(December), 495–502. 10.1016/j.indcrop.2018.12.017

Chen J, Engbersen N, Stefan L, Schmid B, Sun H, & Schöb C (2021). Diversity increases yield but reduces harvest index in crop mixtures. Nature Plants, 7(7), 893–898. 10.1038/s41477-021-00948-4

Costanzo E, Trehin C, & Vandenbussche M (2014). The role of WOX genes in flower development. Annals of Botany, 114(7), 1545–1553. 10.1093/aob/mcu123

Dobin A, Davis CA, Schlesinger F, Drenkow J, Zaleski C, Jha S, Batut P, Chaisson M, and Gingeras TR (2013) “STAR: Ultrafast Universal RNA-Seq Aligner.” Bioinformatics 29, no. 1: 15–21. 10.1093/bioinformatics/bts635.

Doebley JF, Gaut BS, & Smith BD (2006). The Molecular Genetics of Crop Domestication. Cell, 127, 1309–1321. 10.1016.j.cell.2006.12.006

Fujiwara S, Oda A, Yoshida R, Niinuma K, Miyata K, Tomozoe Y, Tajima T, Nakagawa M, Hayashi K, Coupland G & Mizoguchi T (2008) “Circadian Clock Proteins LHY and CCA1 Regulate SVP Protein Accumulation to Control Flowering in Arabidopsis.” Plant Cell 20, no. 11: 2960–71. 10.1105/tpc.108.061531.

Ghidoli M, Pesenti M, Colombo F, Nocito F F, Pilu R, & Araniti F (2023). Camelina sativa (L.) Crantz as a Promising Cover Crop Species with Allelopathic Potential. Agronomy, 13(8). 10.3390/agronomy13082187

Gomez-Cano, F, Carey L, Lucas K, Navarrete TG, Mukundi E, Lundback S, Schnell D, and Grotewold E. (2020) “CamRegBase: A Gene Regulation Database for the Biofuel Crop, Camelina Sativa.” Database 2020 : 1–8. 10.1093/database/baaa075.

Guy SO, Wysocki DJ, Schillinger WF, Chastain TG, Karow RS, Garland-Campbell K, & Burke IC (2014). Camelina: Adaptation and performance of genotypes. Field Crops Research, 155, 224–232. 10.1016/j.fcr.2013.09.002.

Kagale S, Koh C, Nixon J, Bollina V, Clarke WE, Tuteja R, Spillane C, Robinson SJ, Links MG, Clark C et al. (2014) “The Emerging Biofuel Crop Camelina Sativa Retains a Highly Undifferentiated Hexaploid Genome Structure.” Nature Communications 5. 10.1038/ncomms4706.

Kagale S, Nixon J, Khedikar Y, Pasha A, Provart NJ, Clarke WE, Bollina V, Robinson SJ, Coutu C, Wayne D et al. (2016) “The Developmental Transcriptome Atlas of the Biofuel Crop Camelina Sativa.” Plant Journal 88, no. 5 : 879–94. 10.1111/tpj.13302.

King K, Li H, Kang J, & Lu C. (2019). Mapping quantitative trait loci for seed traits in Camelina sativa. Theoretical and Applied Genetics, 132(9), 2567–2577. 10.1007/s00122-019-03371-8

Kreuger F. (The Barbraham Institute). Trim Galore 0.6.10

Ledger S, Strayer C, Ashton F, Kay SA, and Putterill J (2001). “Analysis of the Function of Two Circadian-Regulated CONSTANS-LIKE Genes.” Plant Journal 26(1):15–22.

Lee N, Ozaki Y, Hempton AK, Takagi H, Purusuwashi S, Song YH, Endo M, Kubota A, and Imaizumi T (2023) “The FLOWERING LOCUS T Gene Expression Is Controlled by High-Irradiance Response and External Coincidence Mechanism in Long Days in Arabidopsis.” New Phytologist 239, no. 1 : 208–21. 10.1111/nph.18932.

Li H, Hu X, Lovell JT, Grabowski PP, Mamidi S, Chen C, Amirebrahimi M, Kahanda I, Mummy B, Barry K et al. (2021) “Genetic Dissection of Natural Variation in Oilseed Traits of Camelina by Whole-Genome Resequencing and QTL Mapping.” Plant Genome 14, no. 2 : 1–15. 10.1002/tpg2.20110.

Liang R, Luo C, Liu Y, Hu W, Guo Y, Yu H, Lu T, Chen S, Zhang X, and He X (2023). “Overexpression of Two CONSTANS-like 2 (MiCOL2) Genes from Mango Delays Flowering and Enhances Tolerance to Abiotic Stress in Transgenic Arabidopsis.” Plant Science 327. 10.1016/j.plantsci.2022.111541.

Lin X, Fang C, Liu B, & Kong F (2021). Natural variation and artificial selection of photoperiodic flowering genes and their applications in crop adaptation. ABIOTECH, 2(2), 156–169. 10.1007/s42994-021-00039-0

Liu C, Zhang Q, Zhu H, Cai C, and Li S (2021). “Characterization of Mungbean CONSTANS-LIKE Genes and Functional Analysis of CONSTANS-LIKE 2 in the Regulation of Flowering Time in Arabidopsis.” Frontiers in Plant Science 12(February):1–14. 10.3389/fpls.2021.608603

Liu H, Li Q, & Xing Y (2018). Genes contributing to domestication of rice seed traits and its global expansion. Genes, 9(10). 10.3390/genes9100489

Love MI, Huber W, & Anders S (2014). Moderated estimation of fold change and dispersion for RNA-seq data with DESeq2. Genome Biology, 15(12), 1–21. 10.1186/s13059-014-0550-8

Lu C, & Kang J (2008). Generation of transgenic plants of a potential oilseed crop Camelina sativa by Agrobacterium-mediated transformation. Plant Cell Reports, 27(2), 273–278. 10.1007/s00299-007-0454-0

Lu K, Wei L, Li X, Wang Y, Wu J, Liu M, Zhang C, Chen Z, Xiao Z, Jian H et al. (2019) “Whole-Genome Resequencing Reveals Brassica Napus Origin and Genetic Loci Involved in Its Improvement.” Nature Communications 10, no. 1 : 1–12. 10.1038/s41467-019-09134-9.

Luo Z, Brock J, Dyer JM, Kutchan T, Schachtman D, Augustin M, Ge Y, Fahlgren N, and Abdel-Haleem H (2019) “Genetic Diversity and Population Structure of a Camelina Sativa Spring Panel.” Frontiers in Plant Science 10, no. February : 1–12. 10.3389/fpls.2019.00184.

Malik MR, Tang J, Sharma N, Burkitt C, Ji Y, Mykytyshyn M, Bohmert-Tatarev K, Peoples O, and Snell KD (2018) “Camelina Sativa, an Oilseed at the Nexus between Model System and Commercial Crop.” Plant Cell Reports 37, no. 10 : 1367–81. 10.1007/s00299-018-2308-3.

Müller Niels A., Cris L. Wijnen, Arunkumar Srinivasan, Malgorzata Ryngajllo, Itai Ofner, Tao Lin, Aashish Ranjan, Donnelly West, Julin N. Maloof, Neelima R. Sinha, Sanwen Huang, Dani Zamir, and José M. Jiménez-Gómez (2015) “Domestication Selected for Deceleration of the Circadian Clock in Cultivated Tomato.” Nature Genetics 48(1):89–93.

Nagel DH, & Kay SA (2012). Complexity in the wiring and regulation of plant circadian networks. Current Biology, 22(16). 10.1016/j.cub.2012.07.025

Niwa, Yusuke, Takafumi Yamashino, and Takeshi Mizuno (2009) “The Circadian Clock Regulates the Photoperiodic Response of Hypocotyl Elongation through a Coincidence Mechanism in Arabidopsis Thaliana.” Plant and Cell Physiology 50(4):838–54. DOI: 10.1093/pcp/pcp028

Putri GH, Anders S, Pyl PT, Pimanda JE, & Zanini F (2022). Analyzing high-throughput sequencing data in Python with HTSeq 2.0. Bioinformatics, 38(10), 2943–2945. 10.1093/bioinformatics/btac166

R Core Team (2019). R: A language and environment for statistical computing. R Foundation for Statistical Computing, Vienna, Austria. URL : https://www.R-project.org/

Schiessl S (2020). Regulation and Subfunctionalization of Flowering Time Genes in the Allotetraploid Oil Crop Brassica napus. Frontiers in Plant Science, 11(November). 10.3389/fpls.2020.605155

Schneider CA, Rasband WS, & Eliceiri KW (2012). NIH Image to ImageJ: 25 years of image analysis. Nature Methods, 9(7), 671–675. 10.1038/nmeth.2089

Séguin-Swartz G, Eynck C, Gugel RK, Strelkov SE, Olivier CY, Li JL, Klein-Gebbinck H, Borhan H, Caldwell CD, and Falk KC (2009) “Diseases of Camelina Sativa (False Flax).” Canadian Journal of Plant Pathology 31, no. 4 : 375–86. 10.1080/07060660909507612.

Song JM, Guan Z, Hu J, Guo C, Yang Z, Wang S, Liu D, Wang B, Lu S, Zhou R, et al. (2020). “Eight High-Quality Genomes Reveal Pan-Genome Architecture and Ecotype Differentiation of Brassica Napus.” Nature Plants 6(1):34–45.

Song YH, Shim JS, Kinmonth-Schultz HA, & Imaizumi T (2015). Photoperiodic flowering: Time measurement mechanisms in leaves. Annual Review of Plant Biology, 66, 441–464. 10.1146/annurev-arplant-043014-11555

Soyk S, Müller NA, Park SJ, Schmalenbach I, Jiang K, Hayama R, Zhang L, Van Eck J, Jiménez-Gómez JM, and Lippman ZB (2017) “Variation in the Flowering Gene SELF PRUNING 5G Promotes Day-Neutrality and Early Yield in Tomato.” Nature Genetics 49, no. 1: 162–68. 10.1038/ng.3733.

Takahashi Y, Teshima KM, Yokoi S, Innan H, & Shimamoto K (2009). Variations in Hd1 proteins, Hd3a promoters, and Ehd1 expression levels contribute to diversity of flowering time in cultivated rice. Proceedings of the National Academy of Sciences of the United States of America, 106(11), 4555–4560. 10.1073/pnas.0812092106

Turner A, Beales J, Faure S, Dunford RP, and Laurie DA (2005). “Botany: The Pseudo-Response Regulator Ppd-H1 Provides Adaptation to Photoperiod in Barley.” Science 310 (5750): 1031–34. 10.1126/science.1117619.

Vollmann J, & Eynck C (2015). Camelina as a sustainable oilseed crop: Contributions of plant breeding and genetic engineering. Biotechnology Journal, 10(4), 525–535. 10.1002/biot.201400200

Wang F, Li S, Kong F, Lin X, & Lu S (2023). Altered regulation of flowering expands growth ranges and maximizes yields in major crops. Frontiers in Plant Science, 14(January), 1–14. 10.3389/fpls.2023.1094411

Wang N, Chen B, Xu K, Gao G, Li F, Qiao J, Yan G, Li J, Li H, and Wu X (2016) “Association Mapping of Flowering Time QTLs and Insight into Their Contributions to Rapeseed Growth Habits.” Frontiers in Plant Science 7, no. MAR 2016 (2016): 1–11. 10.3389/fpls.2016.00338.

Wang S. Guo P, Li X, Wu M, Overmyer K, Liu S, & Cui F (2020). Altered redox processes, defense responses, and flowering time are associated with survival of the temperate Camelina sativa under subtropical conditions. Environmental and Experimental Botany, 177 (May), 104132. 10.1016/j.envexpbot.2020.104132

Wei D, Cui Y, He Y, Xiong Q, Qian L, Tong C, Lu G, Ding Y, Li J, Jung C, Qian W (2017). “A Genome-Wide Survey with Different Rapeseed Ecotypes Uncovers Footprints of Domestication and Breeding.” Journal of Experimental Botany 68 (17): 4791–4801. 10.1093/jxb/erx311.

Yuan L, & Li R (2020). Metabolic Engineering a Model Oilseed Camelina sativa for the Sustainable Production of High-Value Designed Oils. Frontiers in Plant Science, 11(February), 1–14. 10.3389/fpls.2020.00011

Zhang H, & Flottmann S (2016). Seed yield of canola (Brassica napus L.) is determined primarily by biomass in a high-yielding environment. Crop and Pasture Science, 67(3–4), 369–380. 10.1071/CP15236

Zhang S, Jiao Z, Liu L, Wang K, Zhong D, Li S, Zhao T, Xu X, and Cui X (2018). “Enhancer-Promoter Interaction of SELF PRUNING 5G Shapes Photoperiod Adaptation.” Plant Physiology 178(4):1631–42.

Zong W, Ren D, Huang M, Sun K, Feng J, Zhao J, Xiao D, Xie W, Liu S, Zhang H et al. (2021). “Strong Photoperiod Sensitivity Is Controlled by Cooperation and Competition among Hd1, Ghd7 and DTH8 in Rice Heading.” New Phytologist 229(3):1635–49.

