## Supporting Information for "Sensitivity to photoperiod is a complex trait in *Camelina sativa*"

Supplemental Figures:

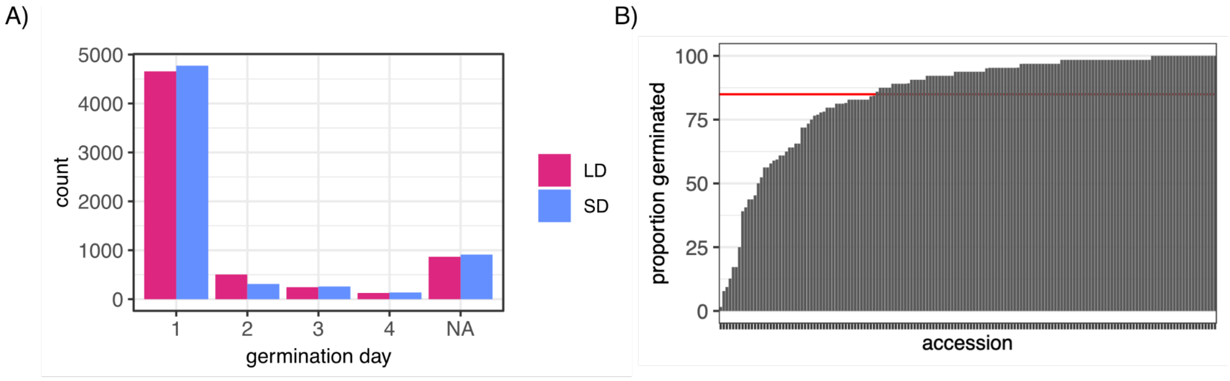


**Supplemental Figure 1. Accessions show a range of both germination day and proportion germinated.** A) Over 70% of total seedlings germinated across all accessions on day 1 (72.77% LD and 74.67% SD ). Of the remaining seeds 7.89% LD and 4.88% germinated on day 2, 3.81% LD and 4.88% SD germinated on day 3 and 1.98% LD  and 2.13% SD germinated on day 4. The remaining seeds 13.54% LD and 14.25% SD did not germinate before the end of this assay (marked with NA on the X-axis). B) The proportion of total seeds that germinated for each accession. The mean (red line) germination rate across all lines was 84.94%.

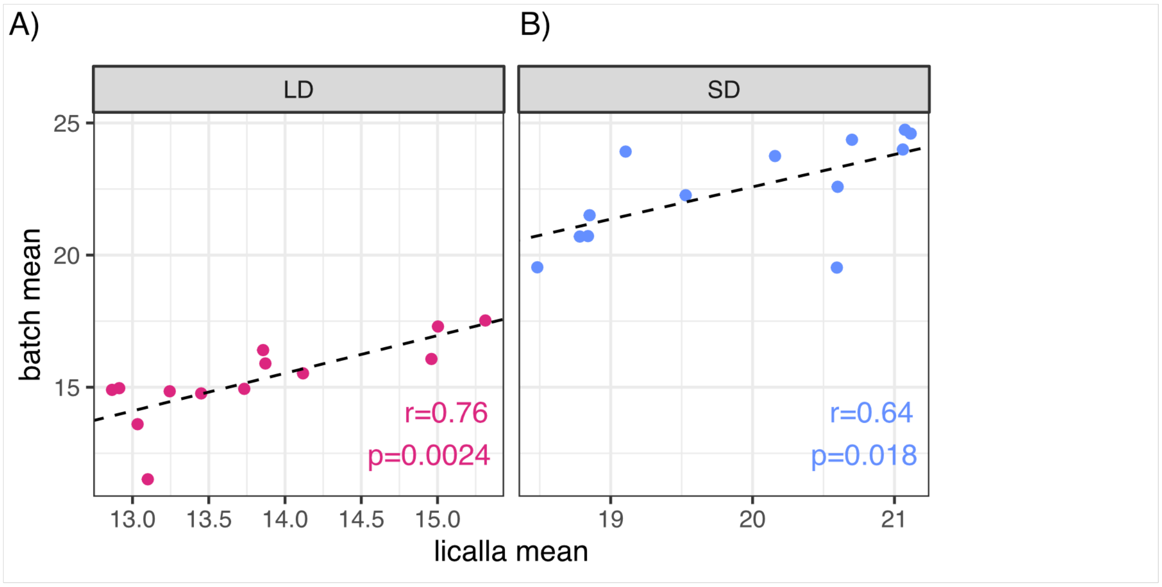


**Supplemental Figure 2. Mean hypocotyl length significantly correlated with Licalla hypocotyl length when stratified by batch.** A) Mean hypocotyl length of tested accessions of each experimental batch (Y-axis) is significantly correlated to mean hypocotyl length of the corresponding Licalla in LD (X-axis) (Pearson correlation : r=0.76, p-value=0.0024). B) This is also true for SD (r=0.64, p-value=0.018).

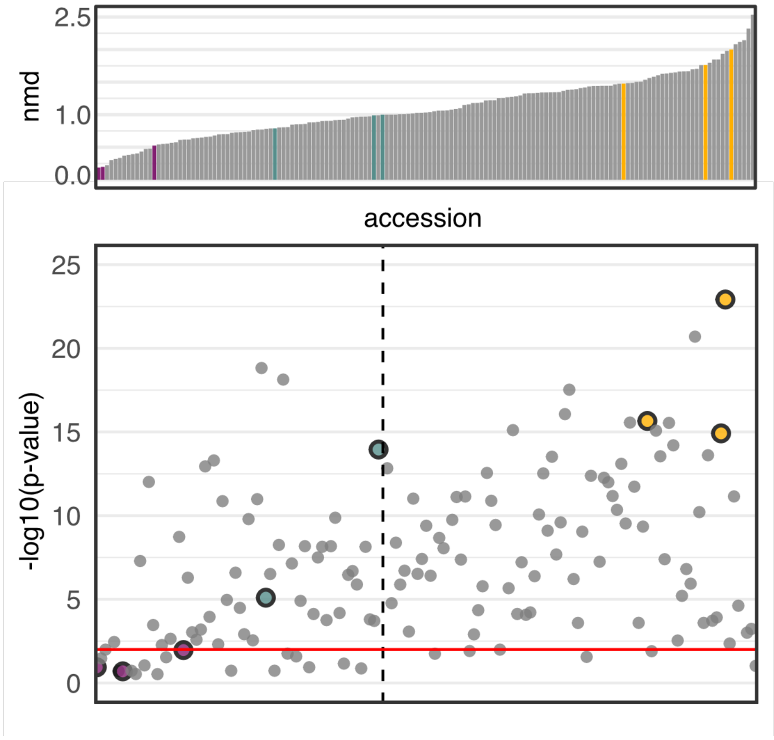


**Supplemental Figure 3. The majority of accessions show significant differences between SD and LD hypocotyl lengths.** Hypocotyl lengths of LD and SD treatments for each accession were tested for significant differences in mean (two-sided t-test). Y-axis shows the -log10(p-value) for each accession (X-axis). Accessions along the X-axis are ordered left to right by increasing NMD score. Red horizontal line is the significance threshold; -log(0.01). The vertical dotted line is the location of Licalla on the X-axis. Low photoperiod-sensitive accessions (CS170, CS129, CS098 ) are marked in purple, high photoperiod-sensitive accessions (CS173, CS096 and CS079) are marked in gold and Licalla (CS002) DH55 are marked in green. Overall, 121 accessions showed significant differences in hypocotyl length between SD and LD treatments and 25 accessions did not. Fifteen accessions did not have enough day-0 seedlings for significance testing.

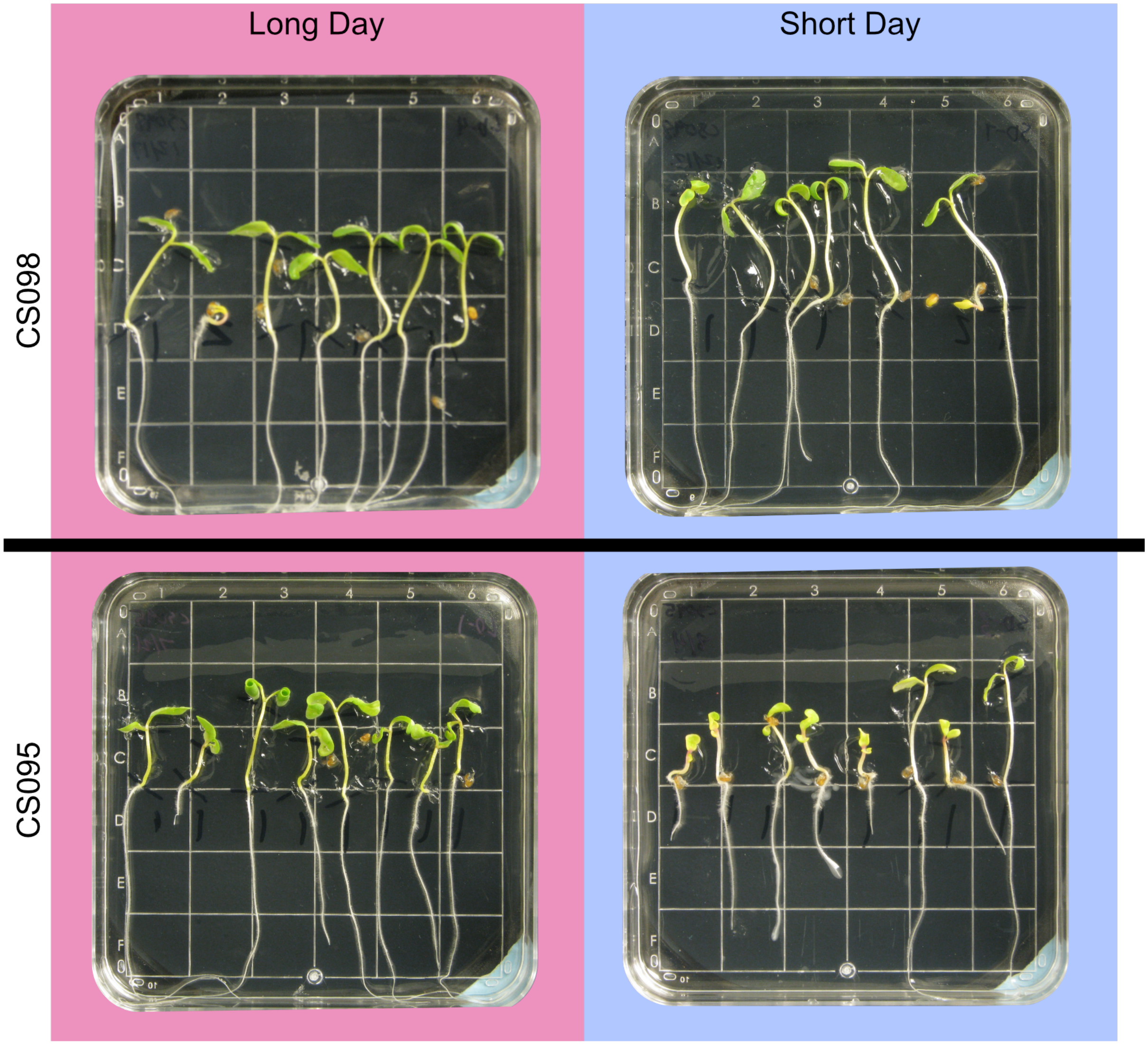


**Supplemental Figure 4. Selecting healthy accessions for validation experiments.**
Representative images of two accessions with low NMD and no significant differences between LD and SD hypocotyl lengths.  Accession CS098 (NMD = -0.18, p-value= 0.022) was considered healthy and selected for retesting. Accession CS095 (NMD = -0.13, p-value= 0.059) was not considered healthy and therefore not selected for validation.

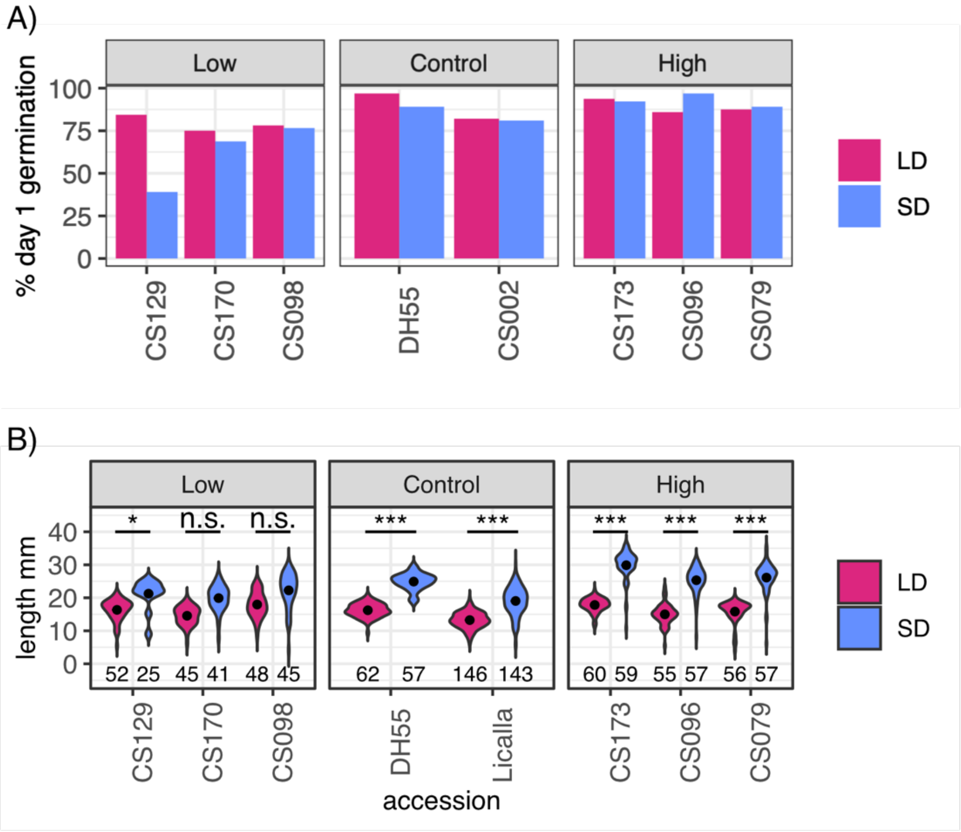


**Supplemental Figure 5: Germination rate and hypocotyl length of LD and SD plants for selected low photoperiod-sensitive, high photoperiod-sensitive and control accessions.** A) The percent of seeds germinated on day one (Y-axis) for each accession (X-axis) was greater than 68% (n=44) for most accessions. Only accession CS129 in the SD condition showed a germination rate of 39% (n=25). B) Hypocotyl length in mm (Y-axis) and accession numbers (X-axis) for SD (blue) and LD (magenta) conditions. In concordance with the initial trial, high photoperiod-sensitive and control accessions showed significant differences between SD and LD growth conditions (two-sided t-test *: $p \leq 0.01$, **: $p \leq 0.001$, ***: $p \leq 0.0001$). One low photoperiod-sensitive accession (CS129) showed a significant difference in hypocotyl length in SD vs LD.

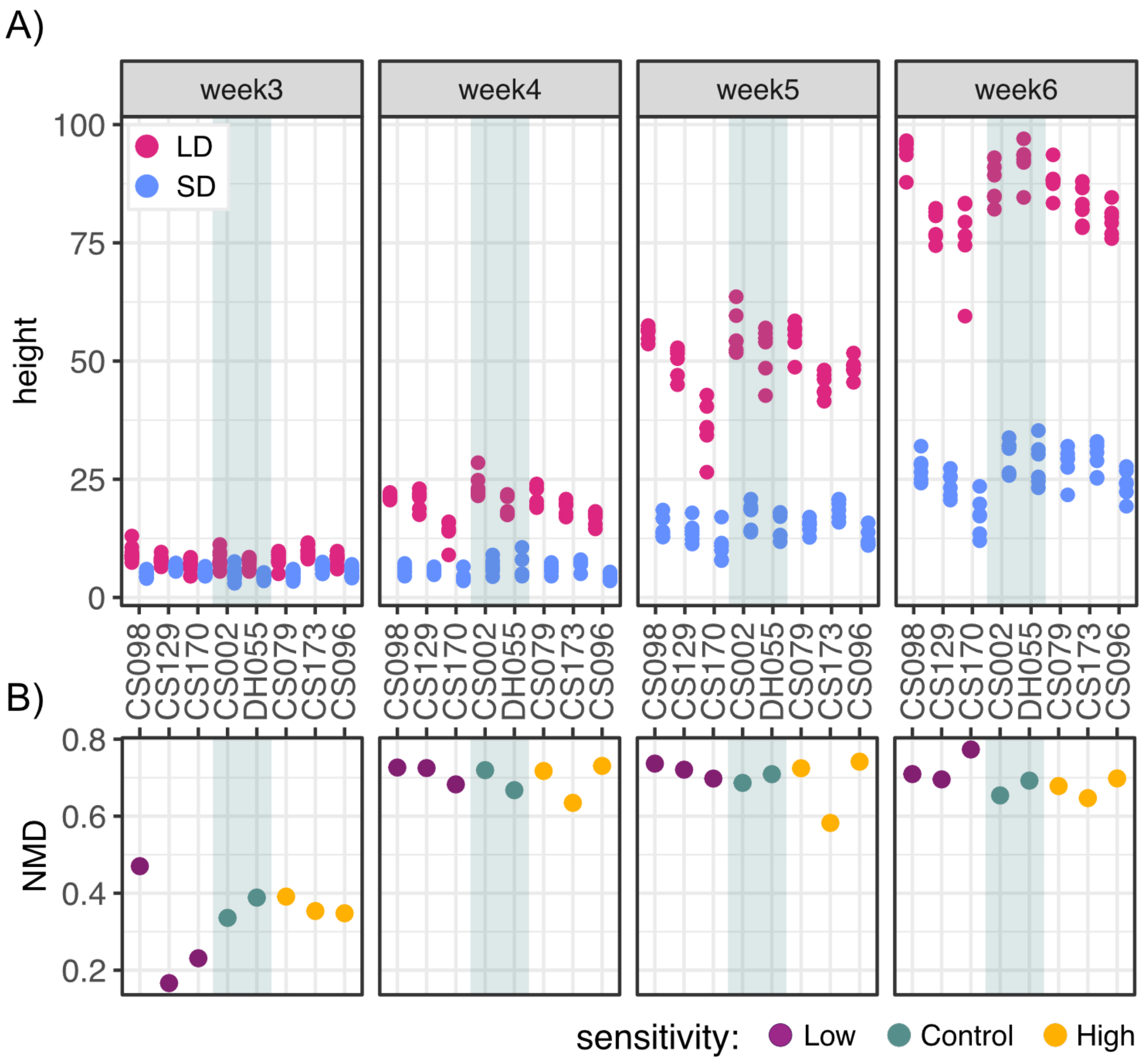


**Supplemental Figure 6. Photoperiod sensitivity in adult plant height varies among accessions over time.** A) Measured height (cm) of soil-grown plants in either LD (magenta) of SD (blue) growth conditions from week 3 to week 6. B) Corresponding normalized mean difference for accessions from week 3 to week 6 (bottom). Low photoperiod-sensitive accessions (CS170, CS129, CS098 ) are marked in purple, high photoperiod-sensitive accessions (CS173, CS096 and CS079) are marked in gold and Licalla (CS002) and DH55 are marked in green.

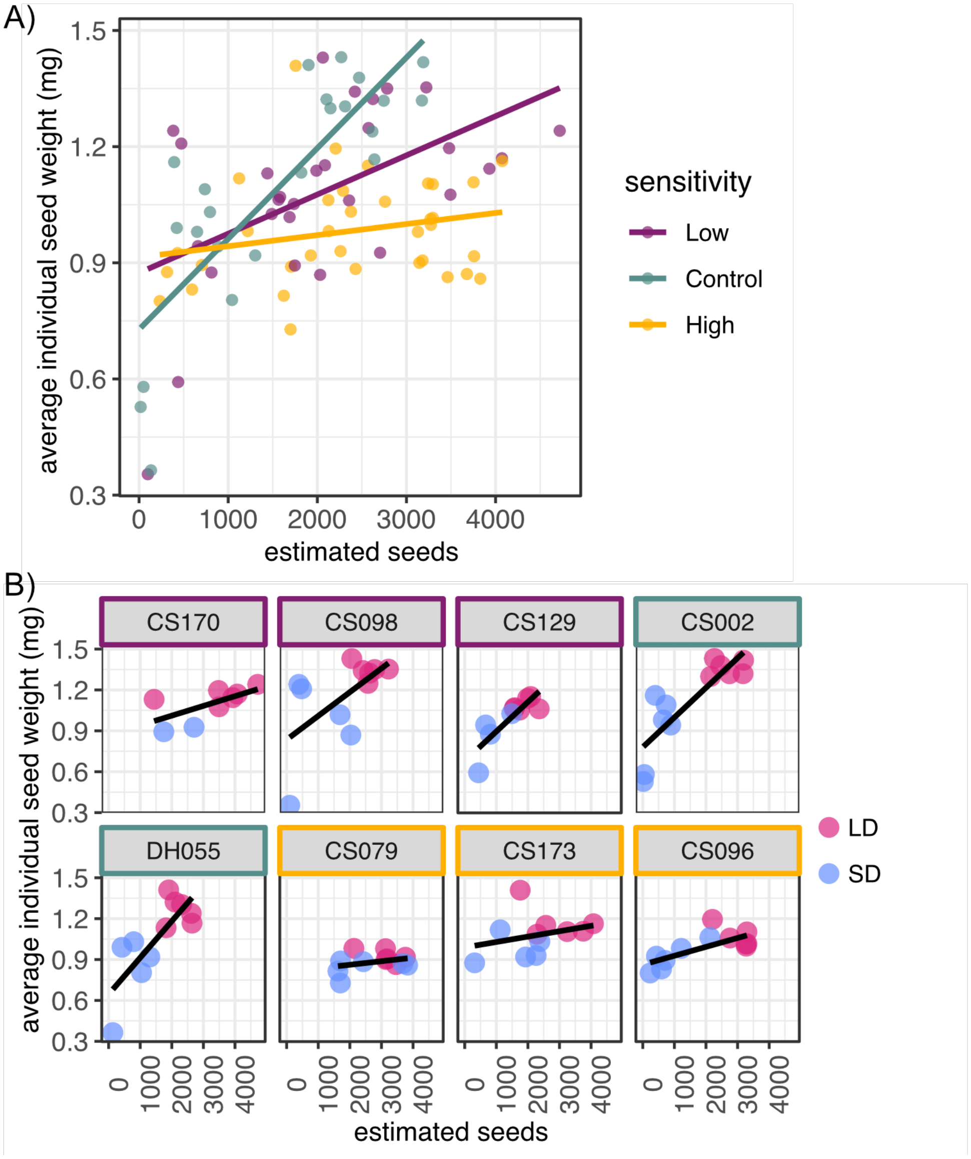


**Supplemental Figure 7. Average individual seed weight and estimated seed number are least correlated in high sensitivity accessions.** A) Average individual seed weight was calculated by weighing 100 seeds from each plant and dividing this weight by 100. The estimated seeds for each plant was calculated by dividing the yield weight by the average individual seed weight. Estimated seed number (X-axis) and the average individual seed weight (Y-axis) are most correlated in the control accessions Licalla and DH55 (Spearman rank correlation, ρ=0.79, p-value= $6.85 \times 10^{-6}$). Similarly, though to a lesser extent, low photoperiod-sensitive accessions were also correlated (Spearman rank correlation, ρ=0.51, p-value=0.005). High photoperiod-sensitive accessions did not show a significant correlation (ρ=0.15, p-value=0.4). Low photoperiod-sensitive accessions (CS170, CS129, CS098 ) are marked in purple, high photoperiod-sensitive accessions (CS173, CS096 and CS079) are marked in gold and Licalla (CS002) DH55 are marked in green. B) The correlation between average individual seed weight and estimated seeds is lost in accession with high sensitivity to photoperiod at the seedling stage.

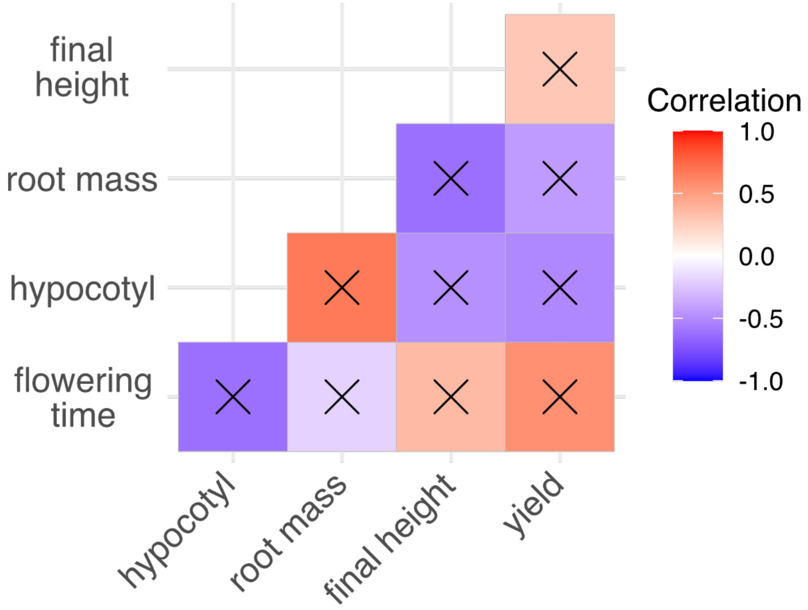


**Supplemental Figure 8: Trait NMD is not significantly correlated across tested Camelina accessions.** NMD values for hypocotyl length (hypocotyl), 20-day root mass (root), final height (measured at day 41), flowering time and seed yield (yield) were pairwise correlated using Spearman's rank correlation. No pairwise correlation was found to be significant, demarcated with “X” in the corresponding pairwise comparison.

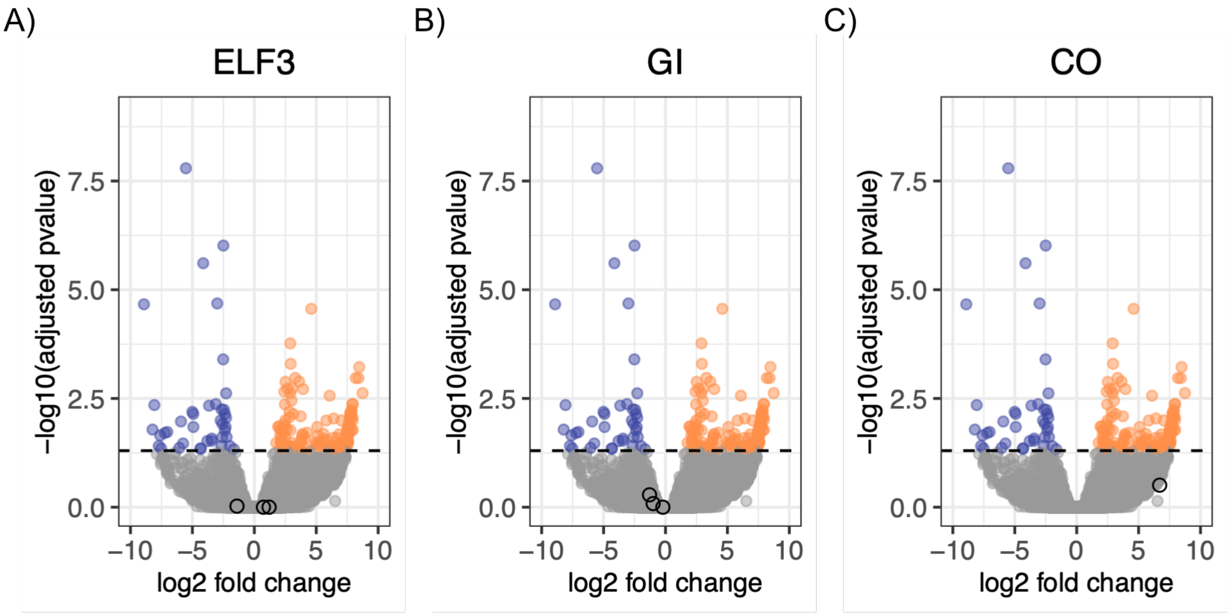


**Supplemental Figure 9: Common flowering time regulators are not differentially expressed between LD and SD conditions.** Expression differences in LD 3-week-old Licalla aerial tissue relative to SD. A total of 40,468 genes were detected with 151 upregulated (orange) and 67 down regulated (blue). Genes that were not found to be differentially expressed are labeled in gray. Syntelogs known to regulate flowering time in *Arabidopsis* that did not show differences in expression are highlighted as black circles. These syntelogs include A) *EARLY FLOWERING 3* (ELF3) B) *GIGANTEA* (GI) and C) *CONSTANS* (CO).

Supplemental Tables :

| Accession number | Accession name | Other names | Biological_status | Origin |
|---|---|---|---|---|
| CS001 | Suneson | MT5 | Cultivar | United States |
| CS002 | Licalla | | Advanced/improved cultivar | Germany |
| CS003 | Kirgizskij | CAM 7 | Advanced/improved cultivar | Kyrgyzstan |
| CS004 | Sortandinskij | CAM 8 | Advanced/improved cultivar | Former Soviet Union |
| CS007 | OmskijMestnyj | CAM 25 | Mutant | Former Soviet Union |
| CS008 | Czestochowska | CAM 28 | Advanced/improved cultivar | Poland |
| CS009 | Volynskaja1 | CAM 29 | Advanced/improved cultivar | Ukraine |
| CS013 | Bronowska1 | CAM 33 | Advanced/improved cultivar | Poland |
| CS015 | Ukrajinskij | CAM 35 | Advanced/improved cultivar | Former Soviet Union |
| CS016 | Wroclawska | CAM 36 | Advanced/improved cultivar | Poland |
| CS018 | CAM38 | CAM 38 | Traditional cultivar/landrace | Austria |
| CS019 | CAM39 | CAM 39 | Traditional cultivar/landrace | Austria |
| CS020 | Lindo | CAM 40 | Advanced/improved cultivar | Germany |
| CS022 | Ukrajinskaja | CAM 45 | Advanced/improved cultivar | Former Soviet Union |

| | | | | |
|---|---|---|---|---|
| CS023 | CAM49 | CAM 49 | | Poland |
| CS024 | CAM52 | CAM 52 | NA | Slovenia |
| CS027 | Kirgizskij2 | CAM 57 | Advanced/improved cultivar | Former Soviet Union |
| CS028 | CAM58 | CAM 58 | Wild | Germany |
| CS030 | Boha1 | CAM 61 | Advanced/improved cultivar | Denmark |
| CS031 | CAME1 | CAM 63 | Advanced/improved cultivar | Swedenden |
| CS033 | CAM66 | CAM 66 | Advanced/improved cultivar | Germany |
| CS034 | Hoga1 | CAM 67 | Advanced/improved cultivar | Denmark |
| CS035 | Svalof | CAM 68 | Advanced/improved cultivar | Sweden |
| CS036 | CAM70 | CAM 70 | Wild | Germany |
| CS037 | Vniimk | CAM 72 | Advanced/improved cultivar | Former Soviet Union |
| CS038 | Kirkizska | CAM 73 | Advanced/improved cultivar | Former Soviet Union |
| CS039 | Czenstochwska1 | CAM 74 | Advanced/improved cultivar | Czech Republic |
| CS042 | PRFGL47 | CAM 79 | Breeding/research material | Germany |
| CS043 | PRFGL36 | CAM 80 | Breeding/research material | Germany |
| CS044 | PRFGL40 | CAM 81 | Breeding/research material | Germany |
| CS046 | PRFGL75 | CAM 83 | Breeding/research material | Germany |

| | | | | |
|---|---|---|---|---|
| CS047 | PRFGL42 | CAM 84 | Breeding/research material | Germany |
| CS048 | PRFGL45 | CAM 85 | Breeding/research material | Germany |
| CS049 | PRFGL57 | CAM 86 | Breeding/research material | Germany |
| CS050 | PRFGL49 | CAM 87 | Breeding/research material | Germany |
| CS051 | PRFGL39 | CAM 88 | Breeding/research material | Germany |
| CS052 | PRFGL72 | CAM 89 | Breeding/research material | Germany |
| CS053 | PRFGL54 | CAM 90 | Breeding/research material | Germany |
| CS054 | PRFGL69 | CAM 91 | Breeding/research material | Germany |
| CS055 | PRFGL59 | CAM 92 | Breeding/research material | Germany |
| CS056 | PRFGL21 | CAM 95 | Breeding/research material | Germany |
| CS057 | PRFGL20 | CAM 96 | Breeding/research material | Germany |
| CS058 | PRFGL19 | CAM 98 | Breeding/research material | Germany |
| CS059 | PRFGL10 | CAM 99 | Breeding/research material | Germany |
| CS060 | PRFGL16 | CAM 101 | Breeding/research material | Germany |
| CS062 | PRFGL34 | CAM 104 | Breeding/research material | Germany |

| | | | | |
|---|---|---|---|---|
| CS063 | Bronowska2 | CAM 108 | Breeding/research material | Poland |
| CS064 | Volynskaja2 | CAM 110 | Advanced/improved cultivar | Poland |
| CS065 | Omskaja | CAM 111 | Advanced/improved cultivar | Former Soviet Union |
| CS067 | PRFGL28 | CAM 114 | Breeding/research material | Germany |
| CS070 | WroclawskaAMCSA70 | CAM 120 | Advanced/improved cultivar | Poland |
| CS071 | Czenstochwska2 | CAM 123 | Advanced/improved cultivar | Czech Republic |
| CS072 | PRFGL22 | CAM 124 | Breeding/research material | Germany |
| CS073 | PRFGL32 | CAM 131 | Breeding/research material | Germany |
| CS074 | CalenaAMCSA74 | CAM 134 | Advanced/improved cultivar | Germany |
| CS075 | CAM135 | CAM 135 | Wild | Germany |
| CS076 | Sortadinskij | CAM 136 | Advanced/improved cultivar | Poland |
| CS077 | CAM137 | CAM 137 | Wild | Denmark |
| CS078 | CAM138 | CAM 138 | Wild | NA |
| CS079 | CAM145 | CAM 145 | Wild | Germany |
| CS080 | STAMM13X14A | CAM 146 | Breeder's line | Germany |
| CS081 | STAMM09X13 | CAM 147 | Breeder's line | Germany |
| CS082 | CAM148 | CAM 148 | Wild | Bulgaria |
| CS083 | STAMM11X15 | CAM 149 | Breeder's line | Germany |
| CS084 | STAMM09X14 | CAM 150 | Breeder's line | Germany |

| | | | | |
|---|---|---|---|---|
| CS085 | STAMM07X13B | CAM 151 | Breeder's line | Germany |
| CS086 | STAMM09X11 | CAM 152 | Breeder's line | Germany |
| CS087 | STAMM08X14 | CAM 153 | Breeder's line | Germany |
| CS089 | STAMM08X13 | CAM 155 | Breeder's line | Germany |
| CS092 | STAMM10X13 | CAM 158 | Breeder's line | Germany |
| CS094 | STAMM13X14B | CAM 160 | Breeder's line | Germany |
| CS095 | STAMM10X15 | CAM 161 | Breeder's line | Germany |
| CS096 | STAMM06X13B | CAM 162 | Breeder's line | Germany |
| CS098 | STAMM06X10A | CAM 164 | Breeder's line | Germany |
| CS103 | Vnttmk17 | CAM 169 | Breeding/research material | Former Soviet Union |
| CS104 | CAM170 | CAM 170 | Wild | Poland |
| CS108 | Voronezskij3491 | CAM 174 | Advanced/improved cultivar | NA |
| CS109 | CAM175 | CAM 175 | Wild | Sweden |
| CS114 | STAMM06X13A | CAM 192 | Breeder's line | Germany |
| CS115 | STAMM06X15B | CAM 193 | Breeder's line | Germany |
| CS116 | STAMM06X15A | CAM 194 | Breeder's line | Germany |
| CS118 | STAMM06X10B | CAM 196 | Breeder's line | Germany |
| CS119 | STAMM02X12A | CAM 197 | Breeder's line | Germany |
| CS120 | STAMM06X07B | CAM 198 | Breeder's line | Germany |
| CS121 | STAMM07X11A | CAM 199 | Breeder's line | Germany |
| CS122 | STAMM06X08A | CAM 200 | Breeder's line | Germany |
| CS125 | STAMM02X11C | CAM 203 | Breeder's line | Germany |
| CS126 | STAMM01X13 | CAM 204 | Breeder's line | Germany |

| | | | | |
|---|---|---|---|---|
| CS127 | STAMM06X09 | CAM 205 | Breeder's line | Germany |
| CS129 | STAMM02X09 | CAM 207 | Breeder's line | Germany |
| CS133 | STAMM06X12B | CAM 211 | Breeder's line | Germany |
| CS136 | STAMM04X13 | CAM 214 | Breeder's line | Germany |
| CS137 | CAM216 | CAM 216 | Traditional cultivar/landrace | Former Soviet Union |
| CS138 | CAM218 | CAM 218 | Traditional cultivar/landrace | Former Soviet Union |
| CS141 | Voronezskij3392 | CAM 221 | Advanced/improved cultivar | Former Soviet Union |
| CS142 | Borow1 | CAM 223 | Advanced/improved cultivar | Poland |
| CS143 | Krajova | CAM 225 | Advanced/improved cultivar | NA |
| CS144 | Vnjjmk17 | CAM 226 | Breeding/research material | Former Soviet Union |
| CS146 | PRFGL96 | CAM 229 | Breeding/research material | Poland |
| CS147 | PRFGL88 | CAM 230 | Breeding/research material | Germany |
| CS148 | PRFGL77 | CAM 231 | Breeding/research material | Germany |
| CS150 | PRFGL95 | CAM 233 | Breeding/research material | Germany |
| CS151 | PRFGL76 | CAM 234 | Breeding/research material | Germany |
| CS152 | PRFGL87 | CAM 235 | Breeding/research material | Germany |
| CS157 | PRFGL78 | CAM 241 | Breeding/research material | Germany |

| | | | | |
|---|---|---|---|---|
| CS159 | CAM243 | CAM 243 | | Germany |
| CS160 | STAMM02X12B | CAM 244 | Wild | Former Soviet Union |
| CS163 | STAMM07X13A | CAM 247 | Breeder's line | Germany |
| CS164 | STAMM06X14A | CAM 248 | Breeder's line | Germany |
| CS166 | STAMM05X14A | CAM 251 | Breeder's line | Germany |
| CS167 | STAMM02X10B | CAM 252 | Breeder's line | Germany |
| CS168 | STAMM06X14B | CAM 253 | Breeder's line | Germany |
| CS170 | STAMM02X08B | CAM 255 | Breeder's line | Germany |
| CS171 | STAMM05X14C | CAM 256 | Breeder's line | Germany |
| CS172 | STAMM06X11B | CAM 257 | Breeder's line | Germany |
| CS173 | STAMM06X13C | CAM 258 | Breeder's line | Germany |
| CS174 | CAM259 | CAM 259 | Wild | Bulgaria |
| CS175 | CAM260 | CAM 260 | Wild | Bulgaria |
| CS176 | CAM261 | CAM 261 | Wild | Bulgaria |
| CS177 | CAM262 | CAM 262 | Wild | Romania |
| CS178 | CAM264 | CAM 264 | Wild | Bulgaria |
| CS179 | CAM266 | CAM 266 | Wild | Former Soviet Union |
| CS180 | CAM268 | CAM 268 | Wild | Bulgaria |
| CS181 | CAM269 | CAM 269 | Wild | Great Britain |
| CS183 | STAMM05X14D | CAM 271 | Breeder's line | Germany |
| CS184 | CAM272 | CAM 272 | Wild | Belgium |
| CS189 | Calinka | CAM 277 | NA | Germany |
| CS191 | GE201101 | Ames 31231 | Wild | Georgia |

| | | | | |
|---|---|---|---|---|
| CS192 | GE201105 | Ames 31232 | Wild | Georgia |
| CS193 | Vniimk17 | PI 258366 | Early variety | Former Soviet Union |
| CS194 | Voronezh349 | PI 258367 | Early variety | Former Soviet Union |
| CS195 | No401 | PI 304268 | Breeding line | Sweden |
| CS196 | No402 | PI 304269 | Breeding line | Sweden |
| CS199 | Borow2 | PI 311735 | donated/collected seed | Poland |
| CS202 | CR47665 | PI 633192 | donated/collected seed | Germany |
| CS204 | GiessenNr3 | PI 633194 | donated/collected seed | Germany |
| CS207 | NU52279 | PI 650141 | donated/collected seed | United States |
| CS208 | CS163 | PI 650142 | donated/collected seed | Denmark |
| CS210 | Boha2 | PI 650144 | donated/collected seed | Denmark |
| CS211 | BRSCHW28347 | PI 650145 | donated/collected seed | Germany |
| CS214 | Giessen3 | PI 650148 | donated/collected seed | Germany |
| CS215 | Giessen4 | PI 650149 | donated/collected seed | Germany |
| CS218 | CPSCAM23 | PI 650152 | donated/collected seed | Germany |
| CS219 | CPSCAM10 | PI 650153 | donated/collected seed | Former Soviet Union |
| CS220 | CSSCAM25 | PI 650154 | donated/collected seed | Former Soviet Union |
| CS225 | CSSCAM33 | PI 650159 | donated/collected seed | Poland |
| CS226 | CSSCAM34 | PI 650160 | donated/collected seed | Former Soviet Union |
| CS227 | CSSCAM35 | PI 650161 | donated/collected seed | Former Soviet Union |
| CS228 | CSSCAM36 | PI 650162 | donated/collected seed | Poland |

| | | | | |
|---|---|---|---|---|
| CS229 | CSSCAM37 | PI 650163 | donated/collected seed | Former Soviet Union |
| CS230 | CSSCAM38 | PI 650164 | donated/collected seed | Austria |
| CS231 | CSSCAM7 | PI 650165 | donated/collected seed | Former Soviet Union |
| CS235 | AMES29309 | PI 652885 | donated/collected seed | Slovenia |
| CS236 | AMES29310 | PI 652886 | donated/collected seed | Slovenia |
| CS245 | MT1 | | Breeder's line | United States |
| CS246 | Blaine Creek | MT3 | Breeder's line | United States |
| CS248 | Celine | | Advanced/improved cultivar | France |
| CS252 | MT148 | | Breeder's line | United States |
| CS253 | MT229 | | Breeder's line | United States |
| CS254 | MT402 | | Breeder's line | United States |
| DH55 | | | Sequencing line | Canada |

**Supplemental Table 1: Name and source information for the 161 accessions included in this study**

| Gene | Arabidopsis syntelog | orientation | Sequence |
|---|---|---|---|
| Csa15g026420 | ACT2* | FWD | CCAGTGTTGTTGGTAGGCCA |
| Csa15g026420 | ACT2* | REV | ACCTCTCTTGGATTGTGCTTCG |
| Csa03g051250 | SEC3B* | FWD | CTAACAATATTCATCCCGCTTCTT |
| Csa03g051250 | SEC3B* | REV | TTTTATATGCCCAGTCAACAACAG |

| | | | |
|---|---|---|---|
| Csa17g071550 | Wox4 | FWD | GCGTCACTTCCGCAACTTTT |
| Csa17g071550 | Wox4 | REV | TTGAGTCGGGTTCCACCTTG |
| Csa05g068740 | FT | FWD | AACCGATTGATTGCATACTCTGATT |
| Csa05g068740 | FT | REV | ACCACCGTTCGTTACTCGT |
| Csa03g002180 | LHY | FWD | CTACATGACAGACTTTGAGGCG |
| Csa03g002180 | LHY | REV | GTGGATAAATCTTAAGCCCAGCC |

**Supplemental Table 2: Primer sequences for qPCR.** * Indicates previously published primers.

| Accession number | SD germination | LD germination | total seeds | day1 germination | Percent D1 germination |
|---|---|---|---|---|---|
| CS001 | 24 | 32 | 64 | 49 | 0.875 |
| CS002 | 1213 | 1219 | 2557 | 2255 | 0.95111459 |
| CS003 | 32 | 29 | 64 | 59 | 0.953125 |
| CS004 | 32 | 31 | 64 | 56 | 0.984375 |
| CS007 | 31 | 32 | 64 | 63 | 0.984375 |
| CS008 | 31 | 31 | 64 | 56 | 0.96875 |
| CS009 | 29 | 31 | 64 | 47 | 0.9375 |
| CS013 | 31 | 32 | 64 | 60 | 0.984375 |
| CS015 | 29 | 20 | 64 | 42 | 0.765625 |
| CS016 | 30 | 19 | 63 | 46 | 0.77777778 |
| CS018 | 24 | 29 | 64 | 35 | 0.828125 |
| CS019 | 25 | 33 | 65 | 55 | 0.89230769 |
| CS020 | 32 | 32 | 64 | 59 | 1 |

| | | | | | |
|---|---|---|---|---|---|
| CS022 | 28 | 25 | 64 | 45 | 0.828125 |
| CS023 | 20 | 32 | 64 | 25 | 0.8125 |
| CS024 | 20 | 30 | 65 | 26 | 0.76923077 |
| CS027 | 24 | 28 | 64 | 40 | 0.8125 |
| CS028 | 32 | 31 | 64 | 46 | 0.984375 |
| CS030 | 25 | 26 | 64 | 40 | 0.796875 |
| CS031 | 28 | 13 | 64 | 16 | 0.640625 |
| CS033 | 19 | 20 | 64 | 27 | 0.609375 |
| CS034 | 29 | 27 | 64 | 48 | 0.875 |
| CS035 | 30 | 29 | 64 | 44 | 0.921875 |
| CS036 | 29 | 29 | 64 | 49 | 0.90625 |
| CS037 | 33 | 31 | 65 | 54 | 0.98461539 |
| CS038 | 32 | 30 | 64 | 52 | 0.96875 |
| CS039 | 32 | 31 | 64 | 60 | 0.984375 |
| CS042 | 32 | 32 | 64 | 47 | 1 |
| CS043 | 29 | 28 | 64 | 51 | 0.890625 |
| CS044 | 31 | 32 | 64 | 52 | 0.984375 |
| CS046 | 31 | 28 | 64 | 44 | 0.921875 |
| CS047 | 30 | 29 | 64 | 58 | 0.921875 |
| CS048 | 31 | 31 | 64 | 45 | 0.96875 |
| CS049 | 30 | 31 | 64 | 52 | 0.953125 |
| CS050 | 30 | 30 | 64 | 50 | 0.9375 |
| CS051 | 32 | 31 | 64 | 61 | 0.984375 |
| CS052 | 32 | 28 | 64 | 53 | 0.9375 |
| CS053 | 31 | 30 | 64 | 46 | 0.953125 |

| | | | | | |
|---|---|---|---|---|---|
| CS054 | 30 | 29 | 64 | 45 | 0.921875 |
| CS055 | 32 | 31 | 64 | 55 | 0.984375 |
| CS056 | 26 | 27 | 64 | 50 | 0.828125 |
| CS057 | 26 | 31 | 64 | 54 | 0.890625 |
| CS058 | 32 | 32 | 64 | 56 | 1 |
| CS059 | 31 | 32 | 64 | 55 | 0.984375 |
| CS060 | 27 | 25 | 64 | 40 | 0.8125 |
| CS062 | 32 | 31 | 64 | 52 | 0.984375 |
| CS063 | 31 | 32 | 64 | 57 | 0.984375 |
| CS064 | 31 | 32 | 64 | 62 | 0.984375 |
| CS065 | 32 | 31 | 64 | 57 | 0.984375 |
| CS067 | 29 | 29 | 64 | 57 | 0.90625 |
| CS070 | 30 | 28 | 64 | 53 | 0.90625 |
| CS071 | 32 | 31 | 64 | 59 | 0.984375 |
| CS072 | 26 | 21 | 64 | 40 | 0.734375 |
| CS073 | 21 | 32 | 65 | 42 | 0.81538462 |
| CS074 | 20 | 20 | 64 | 34 | 0.625 |
| CS075 | 29 | 31 | 64 | 54 | 0.9375 |
| CS076 | 30 | 32 | 64 | 58 | 0.96875 |
| CS077 | 31 | 31 | 64 | 59 | 0.96875 |
| CS078 | 31 | 32 | 64 | 62 | 0.984375 |
| CS079 | 32 | 27 | 64 | 58 | 0.921875 |
| CS080 | 28 | 28 | 64 | 31 | 0.875 |
| CS081 | 30 | 29 | 64 | 44 | 0.921875 |
| CS082 | 32 | 31 | 64 | 54 | 0.984375 |

| | | | | | |
|---|---|---|---|---|---|
| CS083 | 29 | 30 | 64 | 50 | 0.921875 |
| CS084 | 30 | 30 | 64 | 58 | 0.9375 |
| CS085 | 32 | 32 | 64 | 61 | 1 |
| CS086 | 32 | 32 | 64 | 62 | 1 |
| CS087 | 32 | 31 | 64 | 62 | 0.984375 |
| CS089 | 32 | 31 | 64 | 60 | 0.984375 |
| CS092 | 28 | 29 | 64 | 47 | 0.890625 |
| CS094 | 32 | 32 | 64 | 64 | 1 |
| CS095 | 31 | 32 | 64 | 61 | 0.984375 |
| CS096 | 31 | 30 | 64 | 58 | 0.953125 |
| CS098 | 30 | 31 | 64 | 56 | 0.953125 |
| CS103 | 32 | 32 | 64 | 63 | 1 |
| CS104 | 31 | 30 | 63 | 53 | 0.96825397 |
| CS108 | 31 | 30 | 64 | 61 | 0.953125 |
| CS109 | 30 | 30 | 64 | 47 | 0.9375 |
| CS114 | 32 | 32 | 64 | 54 | 1 |
| CS115 | 25 | 25 | 64 | 37 | 0.78125 |
| CS116 | 27 | 26 | 63 | 46 | 0.84126984 |
| CS118 | 19 | 17 | 64 | 29 | 0.5625 |
| CS119 | 32 | 32 | 64 | 64 | 1 |
| CS120 | 27 | 31 | 64 | 44 | 0.90625 |
| CS121 | 29 | 33 | 65 | 56 | 0.95384615 |
| CS122 | 18 | 24 | 64 | 24 | 0.65625 |
| CS125 | 18 | 11 | 64 | 15 | 0.453125 |
| CS126 | 2 | 6 | 63 | 3 | 0.12698413 |

| | | | | | |
|---|---|---|---|---|---|
| CS127 | 32 | 32 | 64 | 63 | 1 |
| CS129 | 28 | 30 | 64 | 53 | 0.90625 |
| CS133 | 29 | 31 | 64 | 55 | 0.9375 |
| CS136 | 29 | 28 | 64 | 53 | 0.890625 |
| CS137 | 30 | 32 | 64 | 55 | 0.96875 |
| CS138 | 31 | 32 | 64 | 57 | 0.984375 |
| CS141 | 0 | 0 | 64 | NA | 0 |
| CS142 | 5 | 11 | 64 | 5 | 0.25 |
| CS143 | 16 | 16 | 64 | 27 | 0.5 |
| CS144 | 0 | 0 | 64 | NA | 0 |
| CS146 | 24 | 27 | 64 | 23 | 0.796875 |
| CS147 | 28 | 25 | 64 | 51 | 0.828125 |
| CS148 | 26 | 29 | 64 | 46 | 0.859375 |
| CS150 | 30 | 32 | 64 | 18 | 0.96875 |
| CS151 | 29 | 31 | 64 | 43 | 0.9375 |
| CS152 | 32 | 32 | 64 | 63 | 1 |
| CS157 | 32 | 32 | 64 | 60 | 1 |
| CS159 | 33 | 32 | 65 | 65 | 1 |
| CS160 | 32 | 32 | 64 | 63 | 1 |
| CS163 | 32 | 32 | 64 | 62 | 1 |
| CS164 | 32 | 32 | 64 | 57 | 1 |
| CS166 | 16 | 22 | 64 | 6 | 0.59375 |
| CS167 | 31 | 32 | 64 | 51 | 0.984375 |
| CS168 | 32 | 31 | 64 | 55 | 0.984375 |
| CS170 | 31 | 32 | 64 | 59 | 0.984375 |

| | | | | | |
|---|---|---|---|---|---|
| CS171 | 32 | 31 | 64 | 52 | 0.984375 |
| CS172 | 25 | 23 | 64 | 25 | 0.75 |
| CS173 | 32 | 32 | 64 | 62 | 1 |
| CS174 | 27 | 30 | 64 | 52 | 0.890625 |
| CS175 | 30 | 32 | 64 | 47 | 0.96875 |
| CS176 | 32 | 32 | 64 | 56 | 1 |
| CS177 | 32 | 32 | 64 | 57 | 1 |
| CS178 | 31 | 32 | 64 | 56 | 0.984375 |
| CS179 | 32 | 32 | 64 | 56 | 1 |
| CS180 | 32 | 31 | 64 | 59 | 0.984375 |
| CS181 | 32 | 32 | 64 | 61 | 1 |
| CS183 | 4 | 2 | 64 | NA | 0.09375 |
| CS184 | 27 | 26 | 64 | 38 | 0.828125 |
| CS189 | 31 | 31 | 64 | 55 | 0.96875 |
| CS191 | 28 | 31 | 64 | 58 | 0.921875 |
| CS192 | 32 | 31 | 64 | 61 | 0.984375 |
| CS193 | 6 | 5 | 64 | 5 | 0.171875 |
| CS194 | 21 | 12 | 63 | 4 | 0.52380952 |
| CS195 | 18 | 18 | 64 | 20 | 0.5625 |
| CS196 | 20 | 22 | 64 | 17 | 0.65625 |
| CS199 | 14 | 14 | 64 | 16 | 0.4375 |
| CS202 | 28 | 28 | 64 | 47 | 0.875 |
| CS204 | 29 | 30 | 64 | 49 | 0.921875 |
| CS207 | 31 | 31 | 64 | 55 | 0.96875 |
| CS208 | 30 | 31 | 64 | 46 | 0.953125 |

| | | | | | |
|---|---|---|---|---|---|
| CS210 | 30 | 31 | 64 | 57 | 0.953125 |
| CS211 | 25 | 26 | 64 | 45 | 0.796875 |
| CS214 | 31 | 31 | 64 | 39 | 0.96875 |
| CS215 | 31 | 32 | 64 | 49 | 0.984375 |
| CS218 | 22 | 24 | 64 | 29 | 0.71875 |
| CS219 | 9 | 24 | 56 | 24 | 0.58928571 |
| CS220 | 19 | 18 | 64 | 30 | 0.578125 |
| CS225 | 3 | 2 | 64 | 1 | 0.078125 |
| CS226 | 16 | 10 | 64 | 15 | 0.40625 |
| CS227 | 0 | 1 | 64 | NA | 0.015625 |
| CS228 | 31 | 31 | 64 | 59 | 0.96875 |
| CS229 | 26 | 27 | 64 | 49 | 0.828125 |
| CS230 | 23 | 18 | 64 | 34 | 0.640625 |
| CS231 | 21 | 25 | 64 | 30 | 0.71875 |
| CS235 | 9 | 19 | 64 | 12 | 0.4375 |
| CS236 | 21 | 18 | 64 | 28 | 0.609375 |
| CS245 | 30 | 30 | 64 | 39 | 0.9375 |
| CS246 | 5 | 6 | 64 | 8 | 0.171875 |
| CS248 | 31 | 30 | 64 | 51 | 0.953125 |
| CS252 | 26 | 28 | 64 | 44 | 0.84375 |
| CS253 | 27 | 26 | 64 | 30 | 0.828125 |
| CS254 | 13 | 12 | 64 | 19 | 0.390625 |
| DH55 | 29 | 31 | 64 | 57 | 0.9375 |

**Supplemental Table 3: Germination day**
